# Protamine 2 Deficiency Results In Septin 12 Abnormalities

**DOI:** 10.1101/2024.05.28.596175

**Authors:** Ondrej Sanovec, Michaela Frolikova, Veronika Kraus, Jana Vondrakova, Maryam Qasemi, Daniela Spevakova, Ondrej Simonik, Lindsay Moritz, Drew Lewis Caswell, Frantisek Liska, Lukas Ded, Jiri Cerny, Tomer Avidor-Reiss, Saher Sue Hammoud, Hubert Schorle, Pavla Postlerova, Klaus Steger, Katerina Komrskova

**Affiliations:** Laboratory of Reproductive Biology, Institute of Biotechnology, Czech Academy of Sciences, BIOCEV, Vestec, Czech Republic; Department of Physiology, Faculty of Science, Charles University, Prague, Czech Republic; Department of Human Genetics, University of Michigan, Ann Arbor, MI, United States; Department of Biological Sciences, College of Natural Sciences and Mathematics, University of Toledo, Toledo, OH, United States; Institute of Biology and Medical Genetics, First Faculty of Medicine, Charles University and General University Hospital in Prague, Prague 2, Czech Republic; Laboratory of Structural Bioinformatics of Proteins, Institute of Biotechnology Czech Academy of Sciences, BIOCEV, Vestec, Czech Republic; Department of Urology, College of Medicine and Life Sciences, University of Toledo, Toledo, OH, United States; Department of Developmental Pathology, Institute of Pathology, University Hospital Bonn, Bonn, Germany; Clinic of Urology, Paediatric Urology and Andrology, Molecular Andrology, Justus Liebig University of Giessen, Giessen, Germany; Department of Zoology, Faculty of Science, Charles University, Prague, Czech Republic

## Abstract

There is a well-established link between abnormal sperm chromatin states and poor motility, however, how these two processes are interdependent is unknown. Here, we identified a possible mechanistic insight by showing that Protamine 2, a nuclear DNA packaging protein in sperm, directly interacts with cytoskeletal protein Septin 12, which is associated with sperm motility. Septin 12 has several isoforms, and we show, that in the *Prm2^-/-^*sperm, the short one (Mw 36 kDa) is mislocalized, while two long isoforms (Mw 40 and 41 kDa) are unexpectedly lost in *Prm2^-/-^* sperm chromatin-bound protein fractions. Septin 12 co-immunoprecipitated with Protamine 2 in the testicular cell lysate of WT mice and with Lamin B1/B2/B3 in co-transfected HEK cells despite we did not observe changes in Lamin B2/B3 protein or SUN4 expression in *Prm2^-/-^*testes. Furthermore, the *Prm2^-/-^* sperm have on average a smaller sperm nucleus and aberrant acrosome biogenesis. In humans, patients with low sperm motility (asthenozoospermia) have imbalanced histone– protamine 1/2 ratio and modified levels of cytoskeletal proteins. We detected retained Septin 12 isoforms (Mw 40 and 41 kDa) in the sperm membrane, chromatin-bound and tubulin/mitochondria protein fractions, which was not true for healthy normozoospermic men. In conclusion, our findings expand the current knowledge regarding the connection between Protamine 2 and Septin 12 expression and localization, resulting in low sperm motility and morphological abnormalities.

## Introduction

Spermiogenesis is a complex process essential for transforming round spermatids into mature sperm. A crucial step in this process is the histone–protamine exchange, which plays an important role in DNA hypercondensation. This exchange is initiated in elongating spermatids and is completed during epididymal maturation. During spermiogenesis, spermatids reduce their size by releasing most organelles and residual cytoplasm in the form of the cytoplasmic droplet (Cooper, 2005), and consequently, by histone–protamine exchange in the nucleus, the highly condensed DNA structure is formed. This DNA hypercondensation is secured by protamines, small arginine-rich proteins, which bind to the minor as well as major grooves of the DNA, resulting in about a 6-fold increase in DNA compaction compared to histones-loaded DNA (Coelingh et al., 1969, Balhorn, 1982, Ward and Coffey, 1991, Mukherjee et al., 2021). This extensive compaction is not only essential for the efficient packaging of DNA but also to protect sperm DNA during the passage through the female reproductive tract. Notably, the replacement of histones by protamines is incomplete, and histone retention in mature sperm varies significantly among species, from 15 % in humans to 1 % in mice (Gatewood et al., 1987, Brunner et al., 2014, Hammoud et al., 2009, Erkek et al., 2013, Hisano et al., 2013).

There are two known types of protamines, protamine 1 and protamine 2, in mammals. While Protamine 1 is present in all mammals, Protamine 2 protein is lost in some species (Ammer et al., 1986). Humans have a 1:1 ratio of P1:P2, whereas in mice, Protamine 2 is more abundant, comprising about 65% of the total Protamine level (Corzett et al., 2002). On the other hand, in mammals such as bulls or boars, Protamine 2 is absent due to the mutation in the promoter region (Maier et al., 1990), and in these species, DNA packaging is organized solely by Prm1. It was shown in humans and could be true for other species, that the protamine ratio is important for sperm fertilizing ability (Schagdarsurengin et al., 2012) and that the relative proportion of sperm Protamine 1 to Protamine 2 mRNA can serve as a valuable clinical parameter to estimate men’s fertility and fertilizing capacity in an assisted reproductive techniques such as IVF (*in vitro* fertilization) or ICSI (intracytoplasmic sperm injection) (Rogenhofer et al., 2013). Moreover, a protamine mRNA ratio likely affects the fertilization ability of sperm and plays a significant role in the initiation of gene expression in the early embryo (Rogenhofer et al., 2017). The negative impact of an aberrant protamine mRNA ratio in embryonic development may be due to changes in chromatin composition and abnormal histone retention (Rogenhofer et al., 2017, Schagdarsurengin and Steger, 2016).

Based on the evidence that the histone–protamine ratio impacts sperm fertilization ability in humans, a protamine 2 knock-out mouse strain (*Prm2^-/-^*) was established (Schneider et al., 2016) to investigate the consequence of loss of protamine 2. Interestingly, *Prm2^+/-^* males are fertile, and sperm retain physiologically normal head morphology and motility, while *Prm2^-/-^* males are sterile, and their sperm display severe phenotypical abnormalities, including fragmented DNA, detached acrosomes at the acrosome-nuclear interface, a higher abundance of reactive oxygen species (ROS), damaged plasma membrane, atypically bent flagellum and loss of motility (Schneider et al., 2016, Schneider et al., 2020). Further, the deletion of *Prm2* in mice affected sperm DNA hypercondensation and integrity and resulted in abnormal acrosome anchorage and flagellum bending, both of which are likely linked to the cytoskeleton function (Schneider et al., 2016).

In spermiogenesis, the cytoskeleton plays a crucial role during acrosome formation, when an actin-keratin structure called acroplaxome emerges. In sperm, the acroplaxome is located in the subacrosomal region and is composed of F-actin and Keratin-5 (Kierszenbaum et al., 2003). This structure is essential for anchoring the acrosome to the nucleus as well as to facilitate myosin-Va mediated vesicular transport by contributing to sperm maturation (Kierszenbaum et al., 2004, Dunleavy et al., 2019). The position of the acroplaxome is likely directed by some proteins of the LINC (LInker of Nucleoskeleton and Cytoskeleton) complex and the ability of keratin-5 to bind to the nuclear lamina (Dunleavy et al., 2019). The LINC complex, which facilitates the connection between the cytoskeleton and nucleoskeleton, is composed of SUN (Sad1/UNc84 homology domain-containing) proteins situated in the inner nuclear membrane and KASH (Klarsicht/Anc1/Syne1 homology domain-containing) proteins located in the outer nuclear membrane (reviewed in (Kmonickova et al., 2020)).

Regarding our study, during spermiogenesis, the localization of SUN protein was shown to be affected by testis-specific Septin 12 (Yeh et al., 2019), which is essential for the correct positioning of the testis-specific SUN4-Lamin B1 complex and sperm head shaping. Lamin Bs are type V intermediate filaments and play various roles in spermiogenesis. Lamin B1 is present in all stages of spermatogenesis, and its function is to decrease the mechanical strength of the nuclear envelope during meiotic chromosome movements (Pereira et al., 2019). On the other hand, Lamin B3 is a spermiogenesis-specific protein. Lamin B3 replaces its longer splicing variant, Lamin B2, during spermiogenesis, and it is suggested that Lamin B3 is essential for nuclear reshaping during spermiogenesis by making the nuclear envelope less stable and more flexible (Furukawa and Hotta, 1993, Schütz et al., 2005a, Schütz et al., 2005b). In relevance to above mentioned connection between SUN and Septin 12 (Yeh et al., 2019), the *Prm2^-/-^*phenotype resembles *SUN4^-/-^*and *Sept12^+/-^* mouse models, particularly the loss of tail elongation and abnormal sperm head shaping, and for *Septin12^+/-^* sperm additionally aberrantly formed tails and decreased motility (Lin et al., 2009, Calvi et al., 2015). Septin 12, as a member of the Septin cytoskeleton protein family, has a conserved GTP binding domain and forms filamentous structures (Mostowy and Cossart, 2012). Septin 12 also contributes to the organization of cytoskeletal proteins and can interact with actin and tubulin and mediates their crosstalk (Nakos et al., 2022). Testis-specific Septin 12 is primarily expressed in male postmeiotic germ cells (Lin et al., 2009, Lin et al., 2011b) with six isoforms described in mice (isoform 1 with calculated molecular mass of 36.0 kDa, isoform 2 of 31.8 kDa, isoform 3 of 40.9 kDa, isoform X1 of 41.3 kDa, isoform X2 of 34.6 kDa, and isoform X3 of 33.3 kDa) and five in humans (isoform 1 with calculated molecular mass of 35.2 kDa, isoform 2 and X2 of 40.7 kDa, isoform X1 of 41.6 kDa, isoform X3 of 34.0 kDa) (https://www.ncbi.nlm.nih.gov/protein/). In mice, Septin 12 is localized to the manchette in elongating spermatids, the neck in elongated spermatids, and the annulus, head, and acrosome in maturated sperm (Yeh et al., 2019). In humans, Septin 12 is present in sperm mainly within the annulus and, to a lesser extent, in the midpiece and head (Lin eta al., 2009). Point mutations affecting the GTP-binding region of human Septin 12 were linked to pathozoospermia, and these mutations are known to disrupt the Septin 12 filamentous structure (Kuo et al., 2012, Lin et al., 2011a). Besides, Septin 12 interaction with α- and β-tubulin are thought to influence head and tail development, supported by sperm of *Septin12^+/−^* males, which displayed unphysiologically formed heads and scattered α- and β-tubulin patterns, resulting in bent and disorganized tails results (Kuo et al., 2012).

To extend the knowledge, we aimed to address the molecular mechanism and protein interactions underlying the above-described phenotypes. We studied the interplay between histone–protamine exchange in connection to sperm motility using the Prm2^-/-^ mouse model and compared the results with those of asthenozoospermic men with aberrant protamine 1/2 ratio. Using biochemical and advanced microscopical techniques, we addressed the localization of Septin 12 during spermiogenesis and in mature mouse and human sperm and defined new potential interaction partners such as Protamine 2, Lamin B2, and Lamin B3.

## Results

### *Protamine 2* deletion disrupted DNA packaging, histone abundance, and selected epigenetic modifications in response

DNA Flow cytometric (FCM) analysis of testicular cell populations represents a rapid, sensitive, and quantitative method for evaluating germ cell maturation during spermatogenesis. Staining DNA in testicular cell suspension followed by FCM allows discrimination of cell populations with different ploidy based on their fluorescence intensity (Rodríguez-Casuriaga and Geisinger, 2021). In our study, using this method, we revealed differences in DNA hypercondensation during spermiogenesis between WT and *Prm2^-/-^* (Fig. 1A), specifically a peak, that represents elongated spermatids and sperm containing sub-haploid DNA content in *Prm2^-/-^* testicular cell samples, was absent. The sub-haploid peak in adult mouse testis cell suspensions reflects the transition from histone–complexed to protamine-complexed DNA, as tight chromatin packaging reduces DNA sites available for fluorochrome binding resulting in an apparent sub-haploid DNA content (Zante et al., 1977) (Spanò and Evenson, 1993). The absence of this peak representing sub-haploid cells indicate disruption of DNA hypercondensation in *Prm2^-/-^* germ cells, and signifies that the DNA condensation in *Prm2^-/-^*sperm is likely resembles elongating spermatids in WT.The other main peaks, which represent haploid cells (1C), diploid cells (2C), and tetraploid cells (4C) were detected in both WT and *Prm2^-/-^*mice (n=5).

**Figure 1:**
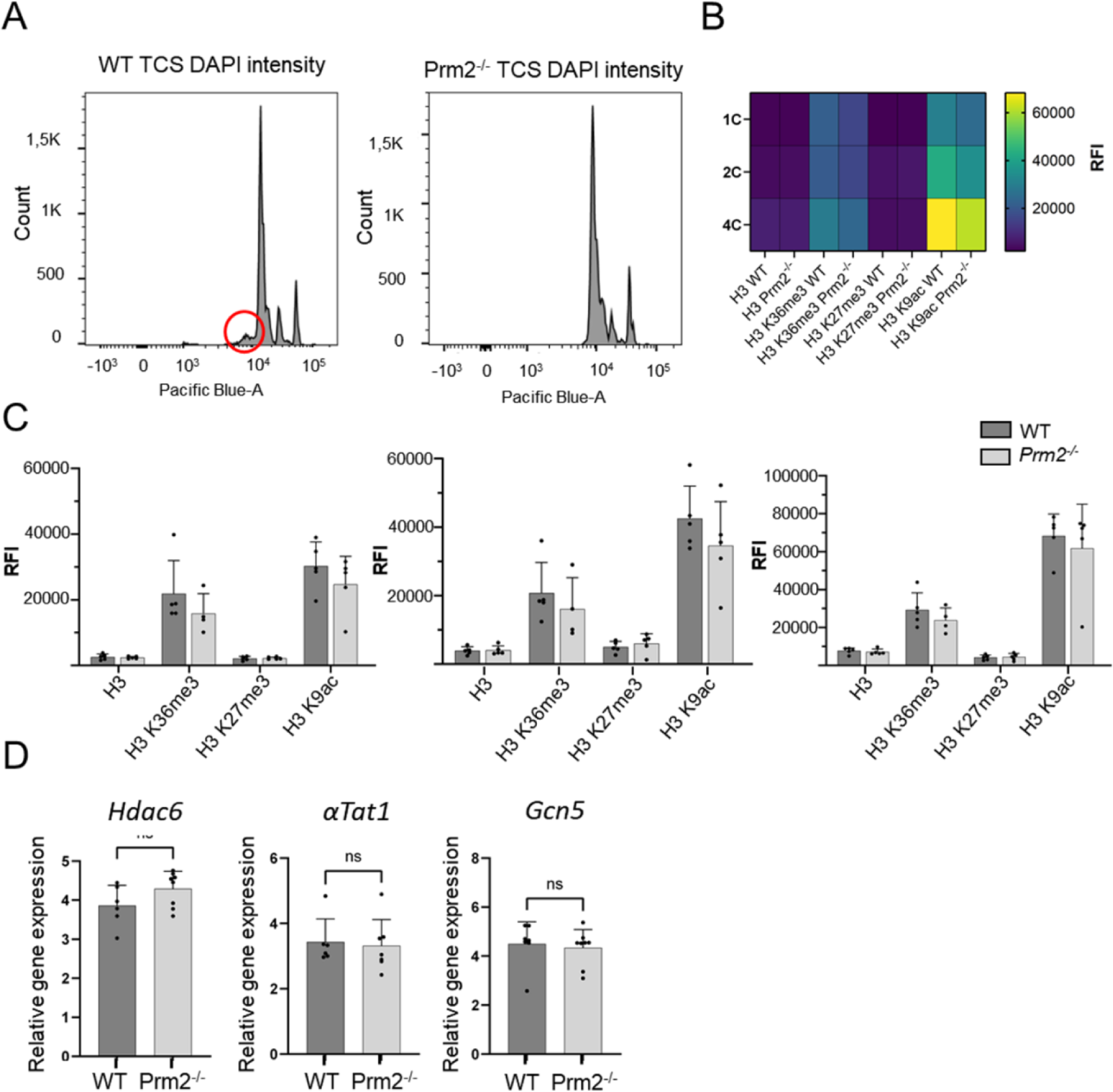
Protamine 2 deletion resulted in histone abundance in testicular cells. **(A)** Distribution of testicular cells based on DAPI intensity (ploidy). The red circle indicates sub-haploid cells present only in WT mouse testis. **(B)** Heat map summarising relative fluorescent intensity (RFI) of all histones stained in all cell populations in both WT and Prm2^-/-^ testicular cells. 1C - haploid cells; 2C - diploid cells; 4C - tetraploid cells. n=5, T-Test, at least 50,000 events were recorded for each measurement. **(C)** Comparison of RFI of individual histones in WT and Prm2^-/-^ in respective cell types. **(D)** qPCR showing no difference in relative gene expression of Hdac6 (Histone deacetylase 6), αTat1 (Alpha-tubulin N-acetyltransferase), and Gcn5 (General control non-depressible 5 also called lysine acetyltransferase 2A) in testicular cells. n=8, Mann-Whitney test.

During the histone–protamine exchange, specific histone modifications (H1, H2A, H2B, H3) play a crucial role (Wang et al., 2019). However, the histone epigenetic modifications in the *Prm2^-/-^* mouse model have remained unexplored. We employed immunofluorescent staining followed by FCM (gating strategy Fig. S1) to investigate potential alterations in epigenetic histone modifications within the *Prm2^-/-^* testicular cells. In the first step, we evaluated the abundance of core histone H3, as well as selected histone modifications, such as H3K27Me3, H3K36Me3, and H3K9Ac in testicular cells (Fig. 1 B, C). The FCM analysis revealed no significant differences in the levels of studied histones and their modifications among the haploid (1C), diploid (2C), and tetraploid (4C) cell populations of both WT and *Prm2^-/-^* mice (n=5) (Fig. 1C). Additionally, qRT-PCR analysis of *Hdac6* (Histone deacetylase 6), *αTat1* (Alpha-tubulin N-acetyltransferase), and *Gcn5 (*General control non-depressible five also called lysine acetyltransferase 2A) in testes did not reveal any difference in the gene expression of these enzymes (Fig. 1D). Our findings indicate that after the depletion of Protamine 2 does not significantly change the abundance of H3 / H3K27Me3, H3K36Me3, H3K9Ac ratio on gene and protein expression levels.

### Sperm morphology is impaired in *Prm2^-/-^* mice

Previous studies characterizing the *Prm2^-/-^* mouse model focused on possible disruption of spermiogenesis and reported mainly sperm damage during the epididymal maturation (Schneider et al., 2016, Schneider et al., 2020). In our study, we focused on the morphology of sperm isolated from the testes, caput, corpus, and cauda of epididymis using Coomassie Brilliant Blue staining (Fig. 2A). Four morphological types of sperm were identified; type 1 – morphologically normal looking; type 2 – tail bent around the head; type 3 – tail partially wrapped around the head; and type 4 – tail completely wrapped around the head (Fig. 2A). Statistical analysis revealed no significant difference in the number of morphologically normal testicular sperm between WT and *Prm2^-/-^* (Fig. 2B, straight dashed line). However, in all epididymal regions, the number of morphologically normal sperm was significantly lower in the *Prm2^-/-^* compared to WT (caput p = 0.0005; corpus and cauda p < 0.0001; Fig. 2B, bend full line). The number of abnormal sperm was also significantly increased in epididymal regions of *Prm2^-/-^* mice compared to WT (caput p = 0.0004; corpus and cauda p < 0.0001). Based on the obtained data and the observation of Schneider and colleagues (2016 and 2020) we concluded that the *Prm2^-/-^* sperm morphology is mainly disturbed during individual stages of epididymal maturation, and the sperm of testicular origin did not display altered morphology compared to WT (Schneider et al., 2016, Schneider et al., 2020).

**Figure 2:**
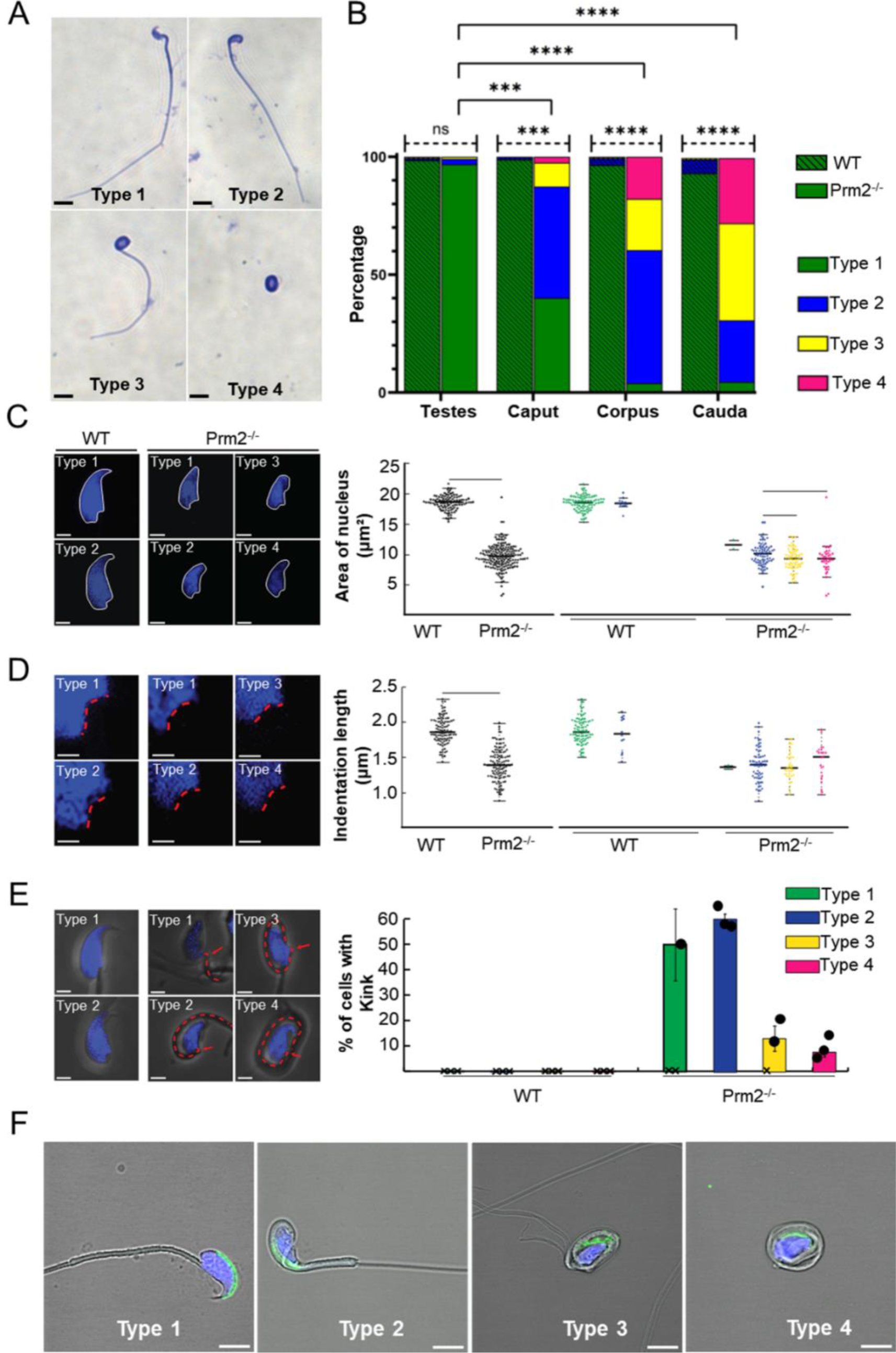
Prm2^-/-^ sperm have pathological morphology during epididymal maturation. **(A)** Representative examples of sperm morphological types present in Prm2^-/-^ cauda epididymis, scale bar = 10 µm. **(B)** Graph shows a significant gradual increase of morphologically abnormal sperm during epididymal maturation. Straight dashed lines indicate the difference in Type 1 sperm between Prm2^-/-^ and WT. The full bent bar represents a decrease of morphologically normal sperm (Type 1) in Prm2^-/-^ in epididymal regions compared to testes. T-test, *** p<0.0005; **** p<0.0001; n=3, at least 200 cells were counted for each individual. **(C)** Area of the nucleus of the cauda epididymis sperm. Left, visualization of sperm head shapes in WT and Prm2^-/-^ stained by DAPI (blue) in the four types we observed, scale bar = 2 µm. The right panel shows nuclei size distribution. The average area of the WT nucleus is 18.47 μm² ± 1.13 (n=106), and that of the Prm2^-/-^ is 9.57 ± 1.99 (n=176), (p<0.0001). The small nuclei size in Prm2^-/-^ is independent of the sperm type. A significant difference between the Prm2^-/-^ Type 2 and Prm2^-/-^ Type 3 (p=0.006) and between Type 2 and Type 4 (p=0.042). **(D)** Nucleus indentations differ in WT and Prm2^-/-^ sperm. Visualization of sperm head shapes in WT and Prm2^-/-^ stained by DAPI (blue) in the four types of sperm. The red line marks the nucleus indentation. The average length of control nucleus indentation is 1.87 μm ± 0.19 (n=101), and that of the Prm2^-/-^ is 1.39 μm ± 0.24 (n=128), (p<0.0001). The nucleus indentation size in Prm2^-/-^ is independent of the sperm type for both control and Prm2^-/-^ sperm. **(E)** Visualization of WT and Prm2^-/-^sperm that present a 90-degree post-neck tail kink (red dashed line). The 90-degree post-neck tail kink frequency was 0 % in WT sperm and 31.8 % in Prm2^-/-^sperm (n=176). The kink was present in 60 % of Type 2 Prm2^-/-^sperm (n=75), 13 % of Type 3 mutant sperm (n=61), and 7.9 % of Type 4 mutant sperm. The 90-degree post-neck tail kink in Prm2 is observed mainly in Types 2 and 3 of mutant sperm. **(F)** Visualization of the acrosomal membrane in Prm2^-/-^ cauda sperm via CD46 (green) immunohistochemistry. Nuclei are visualized in blue (DAPI). Scale bar = 5 µm.

A more in-depth analysis of the caudal sperm revealed a decreased area of the nuclei in *Prm2^-/-^* sperm (Fig. 2C), which is overall in accordance with previously reported findings (Schneider et al., 2020). The detailed measurements delivered more robust data and revealed that the average area of the WT nucleus is 18.47 μm² ± 1.13 (n=106), and that of the *Prm2^-/-^*is 9.57 ± 1.99 (n=176), which is about 50% smaller (p < 0.0001). The small nuclei size in *Prm2^-/-^* is independent of the sperm type. Interestingly, there is a significant difference between the *Prm2^-/-^* sperm Type 2 and Type 3 (p=0.006) and between Type 2 and Type 4 (p=0.042), indicating that Type 2 *Prm2^-/-^* sperm nuclei are larger than the others. In addition, we measured the length of the nuclear indentation of caudal sperm (Fig. 1D) and found out that the average length of the indentation is 1.87 μm ± 0.19 (n= 101) in WT, compared to 1.39 μm ± 0.24 (n= 128) in *Prm2^-/-^*, which is about 25% smaller (p<0.0001). The nucleus indentation size is independent of the sperm type for both WT and *Prm2^-/-^* sperm. There is also a significant difference in the number of sperm 90-degree post-neck tail kink between WT and *Prm2^-/-^* (Fig. 1E). We never observed a 90-degree post-neck tail kink in WT sperm and 32% of *Prm2^-/-^* sperm (n=176). The kink was present in 60% of Type 2 *Prm2^-/-^* sperm (n=75), 13% of Type 3 (n=61), and 7.9% of Type 4 sperm. The 90-degree post-neck tail kink in *Prm2^-/-^* is observed predominantly in Types 2 and 3 of *Prm2^-/-^* sperm.

Further, the acrosome integrity of *Prm2^-/-^* caudal sperm was evaluated. Utilizing immunofluorescent labeling against the CD46 protein as a marker of the acrosomal membranes (Frolikova et al., 2016), we detected the acrosome disintegration in sperm obtained from cauda epididymis (Fig. 2F). Consequently, we examined the acrosomal status of testicular sperm and 3D reconstruction of PNA-stained acrosomes in the individual seminiferous tubule was employed. Using 3D confocal imaging and Imaris software surface rendering, we visualized the abnormal development of acrosomes. Within the sperm head of elongating spermatids, acrosomes of WT could be described as smooth and adjacent to the apical part of the nucleus, whereas in *Prm2^-/-^* the acrosomes were distorted to the side with notable protrusions (Fig. 3). These findings indicate that the abnormal phenotype of *Prm2**^-/-^*** sperm might be established at the beginning or during early stages of spermiogenesis, however, it is fully expanded during epididymal maturation.

**Figure 3:**
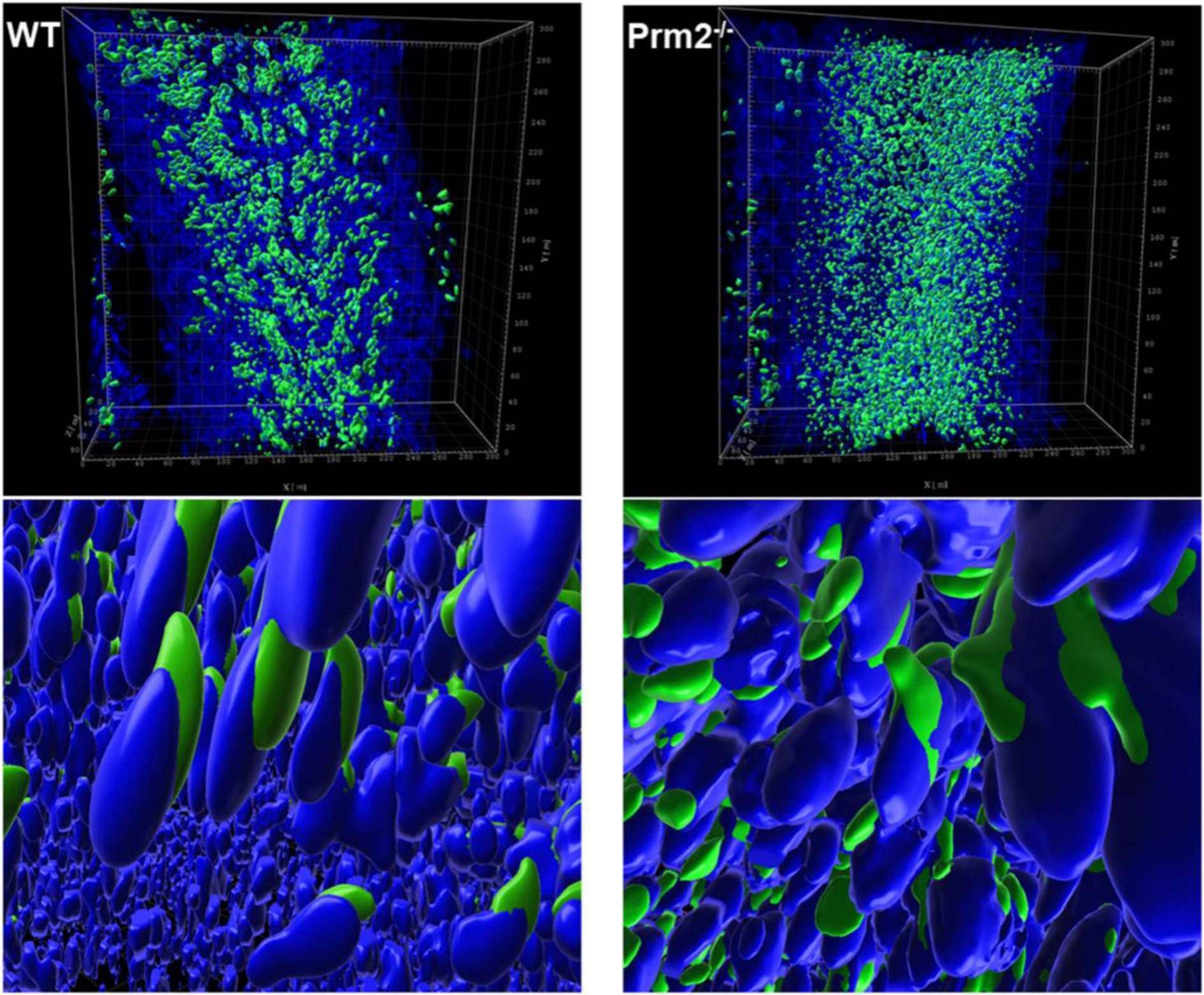
Acrosome formation is aberrant in Prm2^-/-^ seminiferous tubules. 3D visualization of surface-rendered sperm acrosomes (green, PNA-AF488 stained) and nuclei (blue, DAPI stained) in the seminiferous tubule. Surface rendering was postprocessed by Imaris software. The upper panel shows the overall look of half of the seminiferous tubule with surface-rendered acrosomes only. The lower panel shows details of spermatids nuclei and acrosomes in the seminiferous tubule with surface-rendered acrosomes and nuclei.

### Protamine 2 deficiency leads to Septin 12 abnormalities

In spermiogenesis, the cytoskeleton plays a pivotal role in shaping the spermatid head and forming the acrosome, and it is tightly involved in sperm motility. Based on the identification of the abnormal acrosomal shaping in testicular sperm of *Prm2^-/-^* mice coupled with the observation of epididymal sperm motility loss in *Prm2^-/-^*, we evaluated selected cytoskeletal proteins, which could be impaired in *Prm2^-/-^* sperm. The protein abundance of α-tubulin, acetylated α-tubulin at lysine 40 (acK40), β-actin, and Septin 12 in mouse testes and sperm was investigated.

In testicular tissue lysates, western blot analysis, followed by densitometry, was performed (Fig. 4A, B), and the quantification of the obtained data showed no significant difference in the abundance of tubulin and actin proteins between WT and Prm2^-/-^ mice. The presence of two different Septin 12 isoforms in molecular weights of 36 and 40 kDa was detected in both *Prm2^-/-^*and WT testicular tissue. As the antibody used failed to visualize other isoforms, we additionally performed qRT-PCR analysis of *Sept12* in WT and *Prm2^-/-^*testes. Using primers specific for mouse Septin 12 isoforms (1, 2, 3) and a common part of the *Sept12* gene, relative gene expression was calculated with no significant difference between the WT and *Prm2^-/-^* (Fig. 4C).

**Figure 4:**
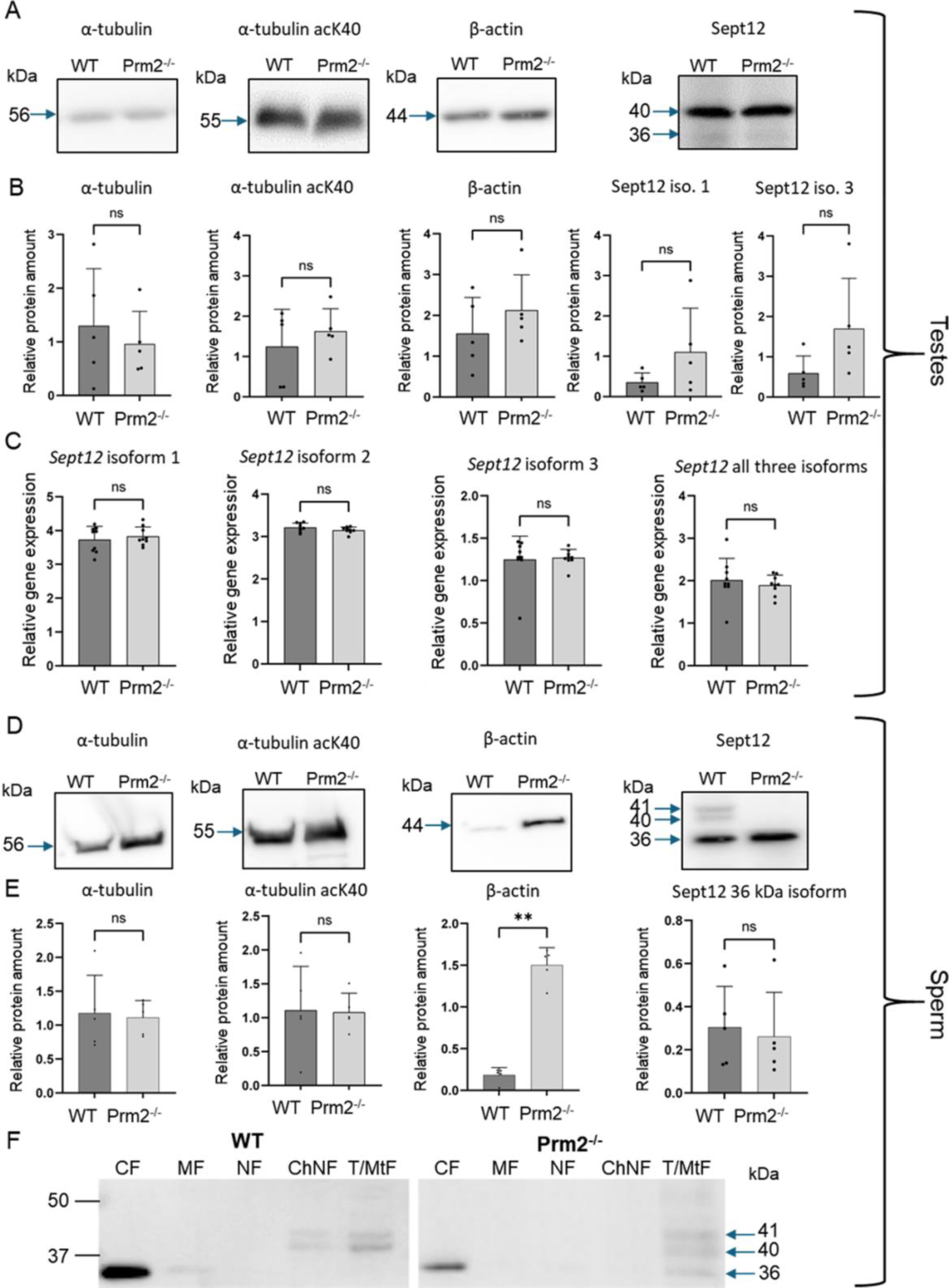
Cytoskeletal protein abundance and gene expression in Prm2^-/-^ sperm and testes is modified for β-Actin, and Septin12 subcellular localization is altered. (A, D) Western blots of specified cytoskeletal proteins in testes (A) or sperm (D) of WT and Prm2^-/-^ mice. Blue arrows indicate the molecular size of the antibody-detected bands. (B, E) A relative protein amount quantification. Band intensities were normalized to the CBB-stained membrane. n=5, Mann-Whitney test, ** p=0.0079. (C) qPCR of total testes lysate showing no difference in relative gene expression of all individual Septin 12 isoforms and total Septin 12. n=8, Mann-Whitney test. (F) Detection of Septin 12 in subcellular fractions of WT and Prm2^-/-^ sperm. CF - Cytoplasmic fraction; MF - Membrane fraction; NF - Nuclear fraction; ChNF - Chromatin-bound nuclear fraction; T/MtF -Tubulin/mitochondria protein fraction. Blue arrows indicate the molecular size of the antibody-detected bands.

In sperm, similarly to the testicular tissue and using the same methodology, the levels of α-tubulin and α-tubulin acK40 remained unchanged (Fig. 4D, E), but we detected a significant increase (p = 0.0079) of β-actin abundance in *Prm2^-/-^* sperm. Importantly, in the sperm lysate from *Prm2*^-/-^ males, only the Septin 12 isoform of 36 kDa was detected while the other two isoforms of 40 and 41 kDa were absent in contrast to WT, where all these isoforms were detected (Fig. 4D). The abundance of 36 kDa isoform in WT and *Prm2^-/-^* showed no significant difference (Fig. 4E).

Using immunofluorescence staining of mature mouse sperm, Septin 12 was shown to be localized in the head, neck, and midpiece of the tail (Lin et al., 2009). To assess potential differences in the localization of individual Septin 12 isoforms across different sperm compartments, we employed western blot analysis of proteins in enriched subcellular fractions from caudal sperm of WT and *Prm2^-/-^*mice (Fig. 4F). Our results revealed that isoform of 36 kDa was present in the cytoplasmic proteins containing fraction (CF) of both WT and *Prm2^-/-^* mouse sperm, as well as in the fraction containing tubulin and mitochondria (T/MtF) of *Prm2^-/-^* sperm. A small amount was also detected in the membrane fraction (MF) of WT, where membrane proteins are enriched. In contrast, both higher-molecular-weight isoforms (40 and 41 kDa) were detected in T/MtF of both WT and *Prm2^-/-^*mouse sperm; however, in WT sperm, both these isoforms were additionally present in chromatin-bound nuclear protein fraction (ChNF) whereas, in *Prm2^-/-^*sperm, was notably absent. The specificity of the Septin 12 antibody was verified on transfected HEK293T/17 cells with mouse Septin 12 plasmid (Fig. S2).

The follow-up detailed analysis of Septin 12 in individual sperm compartments further revealed its presence in the perforatorium and post-acrosomal region of the sperm head, in the connecting piece and midpiece of the tail with notably intense signal observed at the site of the annulus (Fig. 5A) in WT sperm. Although in sperm of Prm2^-/-^ mice, Septin 12 was also identified in both head and tail, it appears that it does not localize into the same compartmental structures as observed in WT sperm. Specifically, the distinct signal detected in the annulus and connecting piece of WT sperm was not observed in *Prm2^-/-^*, nor in the detection of Septin 12 in the perforatorium, which remains unclear. The detailed confocal data provided additional evidence of altered localization of Septin 12 in sperm absence of Protamine 2. To assess whether Septin 12 absence affects its formation in sperm annulus, we employed cryoelectron microscopy (Fig. 5B). There was no visible difference in the annulus shape or localization in Prm2^-/-^ caput sperm to WT; therefore, we analyzed data in FIJI software using Labkit, MorpholibJ and 3DSuite plugins. The applied statistical analysis (Fig. 5C) did not reveal a difference in the annulus area or distance between the centers of the two annular anchoring points (centroid to centroid distance).

**Figure 5:**
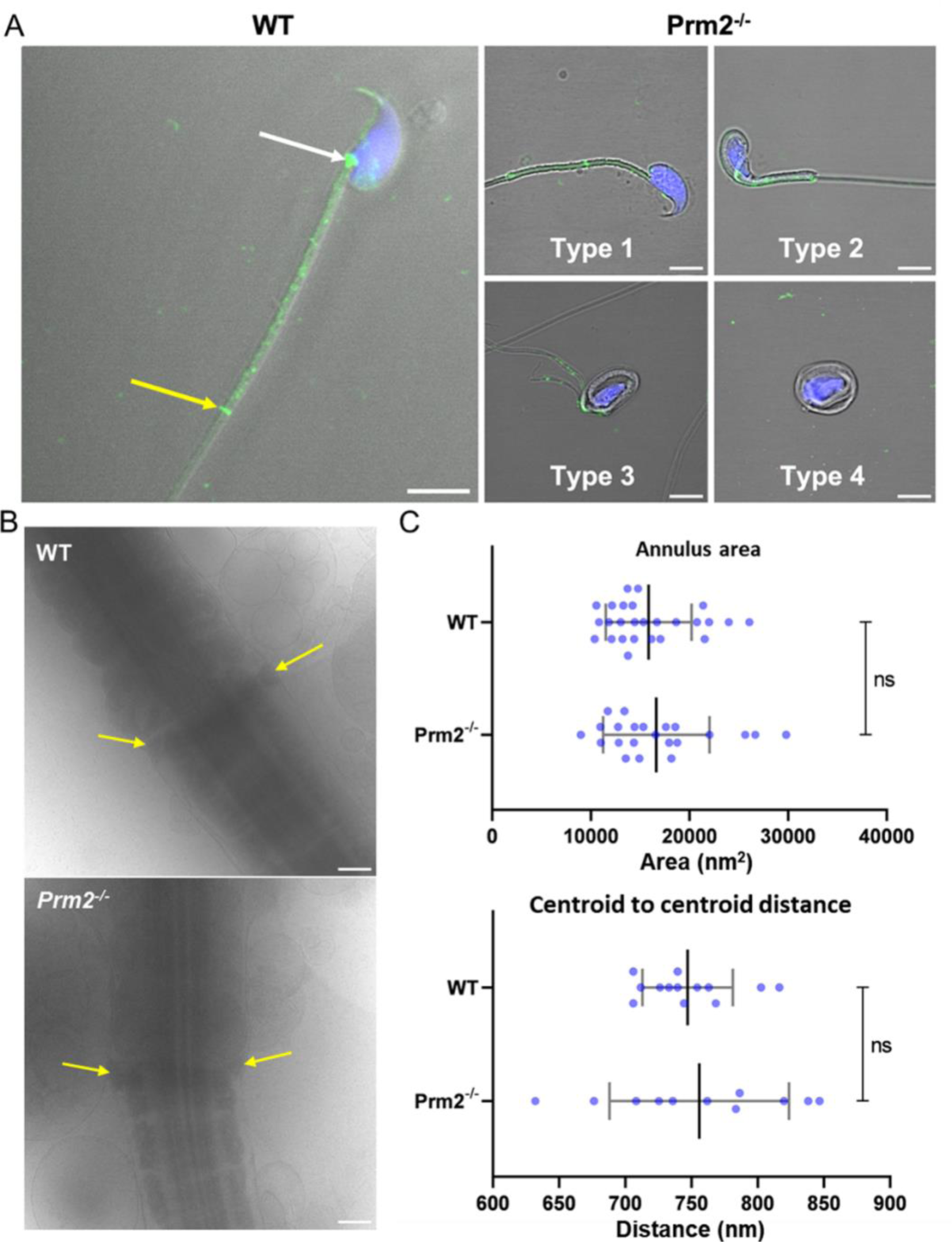
Protamine 2 deletion resulted in aberrant Septin12 localization in epididymal sperm. **(A)** Left panel: A localization of Septin 12 in WT sperm connecting piece (white arrow), annulus (yellow arrow), and midpiece (the area between arrows). Right panel: A mislocalization of Septin 12 in the tail of four morphologically aberrant types of Prm2^-/-^ sperm. Septin 12 (green) and DAPI (Blue), scale bar = 5 µm. **(B)** Annulus (yellow arrows) detail visualized by cryoelectron microscopy. Scale bar = 500 nm. **(C)** Scatterplot visualizing statistical evaluation of mean annulus area and mean of annulus distance from centroid to centroid within one sperm. Mann-Whitney test, n=13 (WT) and 11 (Prm2^-/-^).

There was previously reported interaction between Septin 12 and Pericentriolar material 1 (PCM1) (Yeh et al., 2019), as a component of centriolar satellites, which likely plays a role in the precise localization of various centrosomal proteins and anchoring microtubules to the centrosomes (Dammermann and Merdes, 2002), as demonstrated by yeast two-hybrid assay. Due to the presence of Septin 12 in the sperm connecting piece, where the centrioles are situated, along with a potential link between Septin 12 and centriole function (note that mice have structurally remnant centrioles that maintain centriole proteins (Khanal et al., 2024), we evaluated sperm centriole status in *Prm2^-/-^* mice and implied immunofluorescent staining of Centrosomal Protein 135 (CEP135) (Fig. 6A, Fig. S3). Using the Leica LAS-X software algorithm we analyzed confocal images concluding, that *Prm2^-/-^* sperm centriole remnant is abnormal. Both the mean (Fig. 6B) and maximum (Fig. 6C) intensities of the CEP135 signal are reduced by about 35 % (p = 4.4×10^-6^ and p = 5.8×10^-7^, respectively). Moreover, the *Prm2^-/-^* Cep135-labeled centriole is 10% longer (Fig. 6D) than the WT (p = 0.04). The average length of the control group Cep135-labeled centriole was 0.74 ± 0.20 μm while the length of *Prm2^-/-^*centriole was 0.81 ± 0.15 μm. Interestingly, the average width (Fig. 6E) of control and *Prm2^-/-^* Cep135-labeled centrioles was 0.47 ± 0.09 μm and 0.48 ± 0.10 μm, respectively, showing no significant difference.

**Figure 6:**
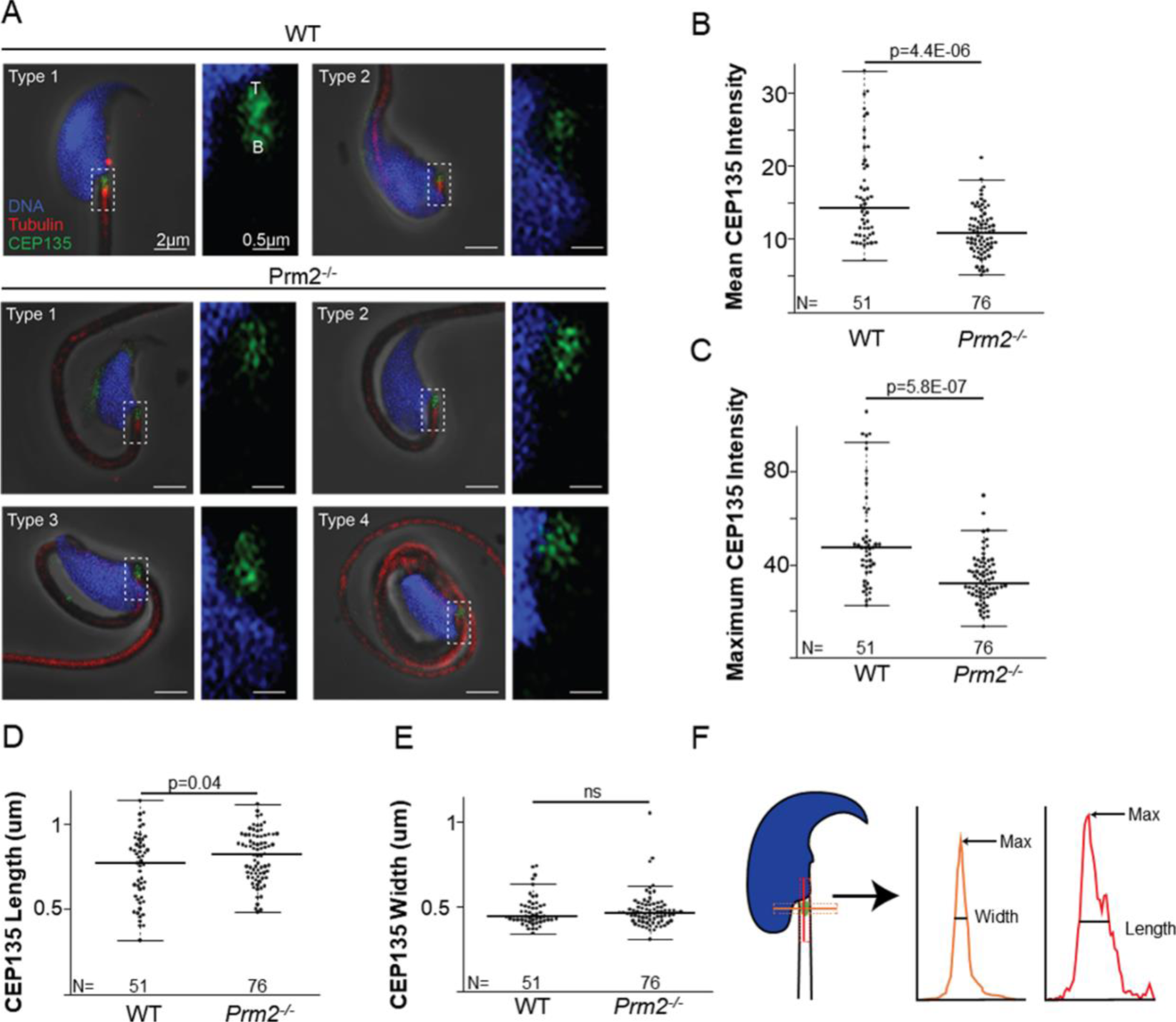
Prm2^-/-^ spermatozoa remnant centrioles are abnormal. **(A)** Examples of WT types 1 and 2 spermatozoa and the four types of Prm2^-/-^ spermatozoa. Each type has two images: the left panel shows a low-magnification view of the sperm head, and the right panel illustrates a high-magnification view focusing on the centriolar staining. CEP135 is shown in green, the tail axoneme in red, and the nucleus in blue. All sperm were oriented with the tip of the centriole (T) at the top of the image and the base of the centriole (B) at the bottom. Scale bar = 2 and 0.5 µm, respectively. **(B)** Graph illustrating CEP135 mean intensity. The wild-type spermatozoa average intensity was 15.92 ± 6.46 (n=51), and the average for the mutant was 11.00 ± 3.13 (n=76), resulting in 30 % reduced labeling in Prm2^-/-^ (p=4.4E-06). The sperm mean intensity was quantified using a Leica LAS-X algorithm. **(C)** Graph representing CEP135 maximum intensity. The average maximum intensity for the wild-type sperm was 51.36 ± 21.07 (n=51), and for the mutant sperm was 33.62 ± 10.22 (n=76). Signal intensity is 35 % reduced in Prm2^-/-^ (p=5.8E-07). **(D)** The CEP135 labeling length is 10% longer (p=0.04) in Prm2^-/-^ than in wild-type (WT). The average CEP135 length of the WT sperm was 0.74 ± 0.20 μm (n=51), and the Prm2^-/-^ was 0.81 ± 0.15 μm (n=76). **(E)** The CEP135 labeling width length is similar (p=0.705) in Prm2^-/-^ and WT. The average CEP135 width of WT sperm was 0.47 ± 0.09 μm (n=51), and of the mutant was 0.48 ± 0.10 μm (n=76). **(F)** Schematic illustration showing CEP135 length and width measurement strategy. Width is shown in orange, and length in red.

### Interaction of Septin 12 with Protamine 2 and Lamin B2/3 intermediate filaments

Based on findings that the lack of Protamine 2 might be responsible for the absence of Septin 12 isoform 3 in *Prm2^-/-^* mouse sperm, we employed a co-immunoprecipitation assay to explore their potential interaction. The Septin 12 – Protamine 2 complex was precipitated from mouse testes lysate using an anti-Protamine 2 antibody. Subsequently, the samples were immunodetected with an anti-Septin 12 antibody (Fig. 7A). Two bands corresponding to the molecular weights of 36 and 41 kDa were observed in the precipitated sample, while it was absent in the IgG control, indicating a possible interaction between Septin 12 and Protamine 2.

**Figure 7:**
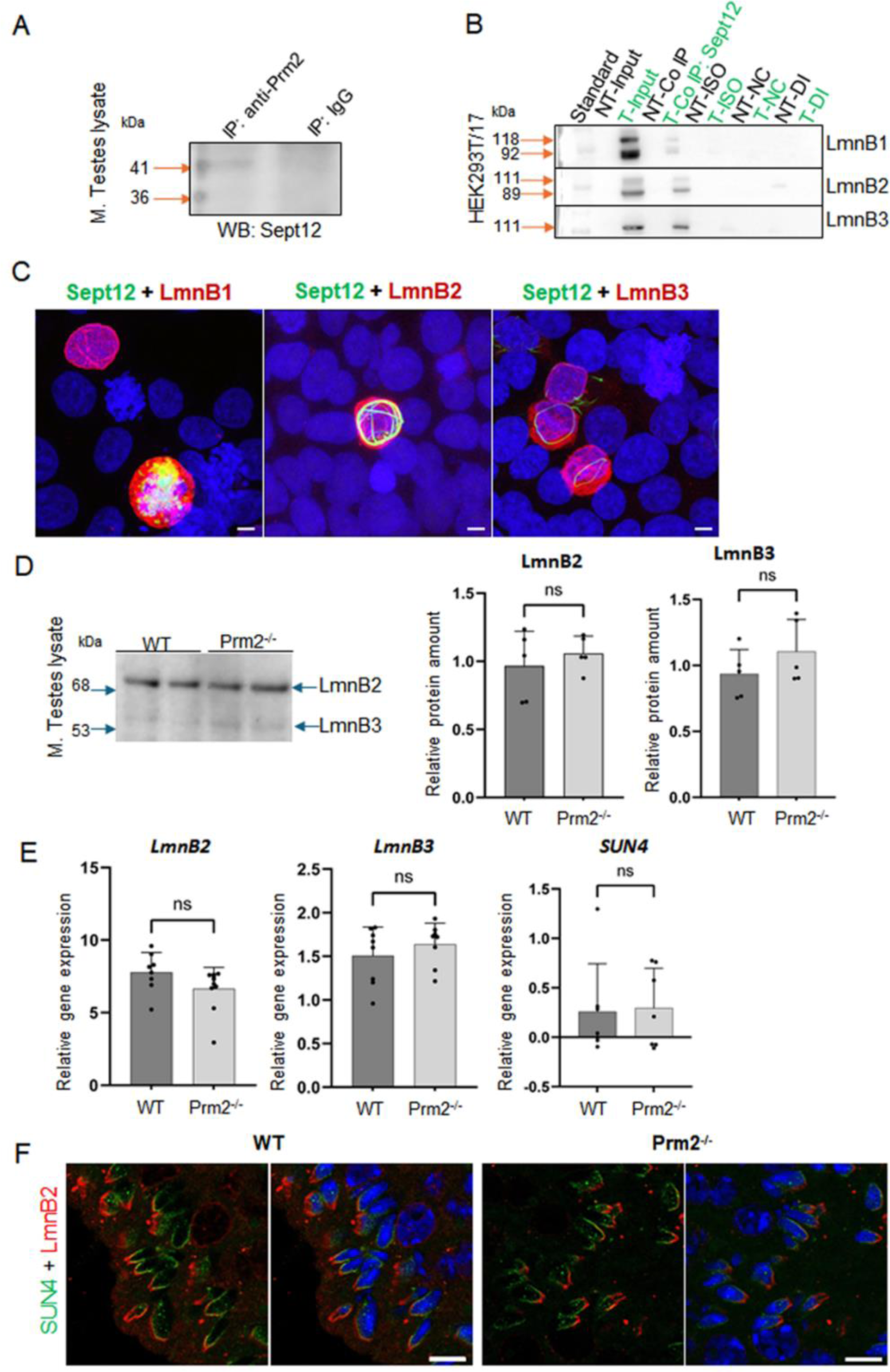
Analysis of Septin 12 interacting proteins reveals positive interaction with Protamine 2 and Lamin B2/3. **(A)** Co-immunoprecipitation of Protamine 2 with Septin 12 in mouse testes lysate. Blue arrows indicate a molecular weight standard; orange arrows indicate a precipitate in predicted molecular weights. Isotype IgG control shows no precipitate. **(B)** Co-immunoprecipitation of transfected HEK cells with mouse Septin 12 and Lamin B1, Lamin B2, or Lamin B3. NT - non-transfected cells; T - Transfected cells; Input-only cell lysate (without Co-IP); Iso - Isotype control; NC - Negative control for Co-IP (without ab); DI - Depleted input (the last wash of Co-IP). **(C)** 3D illustrative images showing positive transfection of HEK cells with Septin 12-GFP (green) and Lamin Bs-Myc (red). Nuclei are stained by DAPI (blue). Scale bar = 5 µm. **(D)** Western blot shows no significant difference in the relative abundance of Lamin B2 and Lamin B3 in mouse testes. Blue arrows indicate the molecular size of the antibody-detected bands. **(E)** qPCR of mouse testes shows no significant difference in relative gene expression of Lamin B2, Lamin B3, and SUN4 in testicular cells of WT and Prm2^-/-^ mice. n=8, Mann-Whitney test. **(F)** Immunohistochemistry of mouse SUN4 (green) and Lamin B2 (red). Scale bar = 10 µm.

**Figure 8:**
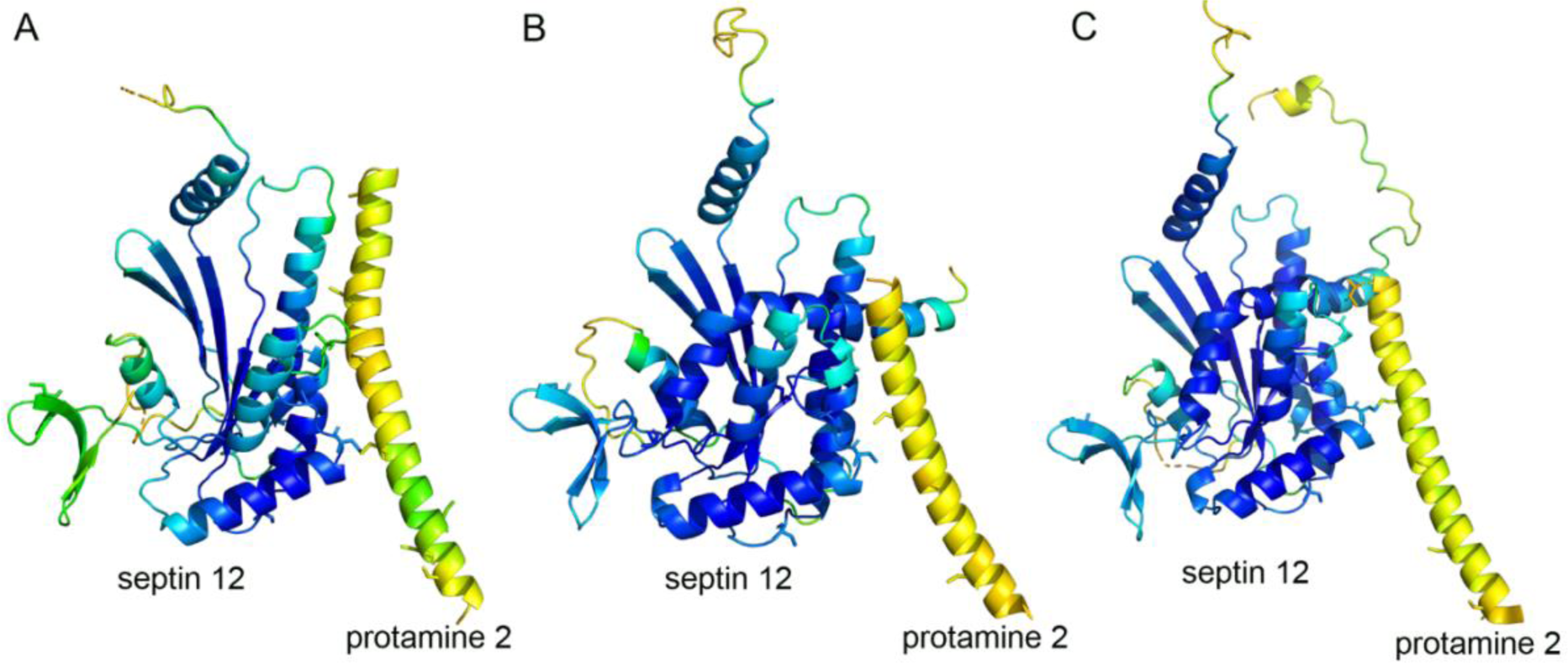
Prediction of Protamine 2/Septin 12 complexes with AlphaFold multimer. The proteins are shown in cartoon representation with Cysteine residues in sticks representation. Residues are color coded according to the transformed pLDDT confidence score using 100-pLDDT with the standard B-factor color scale with cold colors corresponding to high confidence regions. Only residues with pLDDT score higher than 40 are shown. **(A)** Predicted complex of mouse Protamine 2 and Septin 12 isoform 1. **(B)** Predicted complex of mouse Protamine 2 and Septin 12 isoform 3. **(C)** Predicted complex of human Protamine 2 and Septin 12 isoform 2.

Within the sperm nucleus, Septin 12 was shown to interact with the SUN4/Lamin B1 complex (Yeh et al., 2015). In the current study, HEK293T/17 cells were co-transfected with the Septin 12-GFP and LaminBx-Myc plasmids, followed by co-immunoprecipitation assay to investigate the possible interaction of Septin 12 with other Lamin family members – specifically Lamin B2 and Lamin B3. Co-transfection of Septin 12 with Lamin B1 served as a positive control. The Septin 12-Lamin Bx complex was precipitated via GFP-tag on Septin 12 using an anti-GFP antibody, followed by immunodetection using an anti-Myc antibody. Myc-tag labeling revealed the double band corresponding to two isoforms of Lamin B1 (118 kDa, 92 kDa), as well as two bands for Lamin B2 (111 kDa, 89 kDa). Only a single band with a molecular weight of 111 kDa corresponding to Lamin B3 was detected. No immunodetection was observed in the non-transfected cell lysate or the IgG control. (Fig. 7B). Co-transfected HEK293T/17 cells are shown in representative Fig. 7C. These data suggested that Septin 12 interacts with the investigated Lamin B proteins either directly or indirectly within a protein complex.

To inspect the potential influence of lack of *Prm2* on the abundance of Lamin B proteins, we performed a western blot analysis using mouse testes lysates (Fig. 7D). The bands corresponding to molecular weights of Lamin B2 (68 kDa) and Lamin B3 (53 kDa) were detected. However, densitometric analysis revealed no significant change in the levels of Lamin B2 or B3 in the *Prm2^-/-^* testes compared to WT. Furthermore, using qRT-PCR analysis of cDNA from mouse testes, we also did not detect a significant difference between WT and *Prm2^-/-^* and the relative gene expression levels of *LaminB2* and *LaminB3*, as well as S*UN4* (Fig. 7E), suggesting no difference in the regulation of these proteins on expression level. To compare the localization of Lamin B2 and SUN4 in testicular cells, we employed indirect immunohistochemistry of FFPE mouse testes. We were able to detect SUN4 as well as Lamin B2 in postmeiotic cells, in particular in elongating/elongated spermatids (Fig. 7F) in the lateral posterior part of the spermatid head, where the manchette is localized, with absence in the head-to-tail coupling apparatus. These results suggest that the absence of Protamine 2 does not significantly affect Lamin B2/3 and SUN4 expression, protein abundance, and localization within nuclear lamina composition in *Prm2^-/-^* mice testicular cells.

A possible interaction is also predicted by the AlphaFold multimer for Protamine 2 with Septin 12 model complexes. The modeling predicts association of a helical C-terminal Protamine 2 region (residues from about 60 to 100) with the Septin 12 protein. Although the structure of the C-terminal Protamine region is modeled with relatively low confidence (pLDDT between 50 and 60), the potential interaction could be further stabilized by predicted disulfide bridges between Protamine 2 and Septin 12. The potential disulfide bridges involve mouse Protamine 2 Cys88 and Septin 12 isoform 1 Cys217, mouse Protamine 2 Cys70 and Septin 12 isoform 3 Cys217, and human Protamine 2 Cys74 and Septin 12 isoform 2 Cys219 (homologous to the murine Cys217), respectively. The Cysteine residues in predicted complexes are shown in sticks representation in the Fig 9.

**Figure 9:**
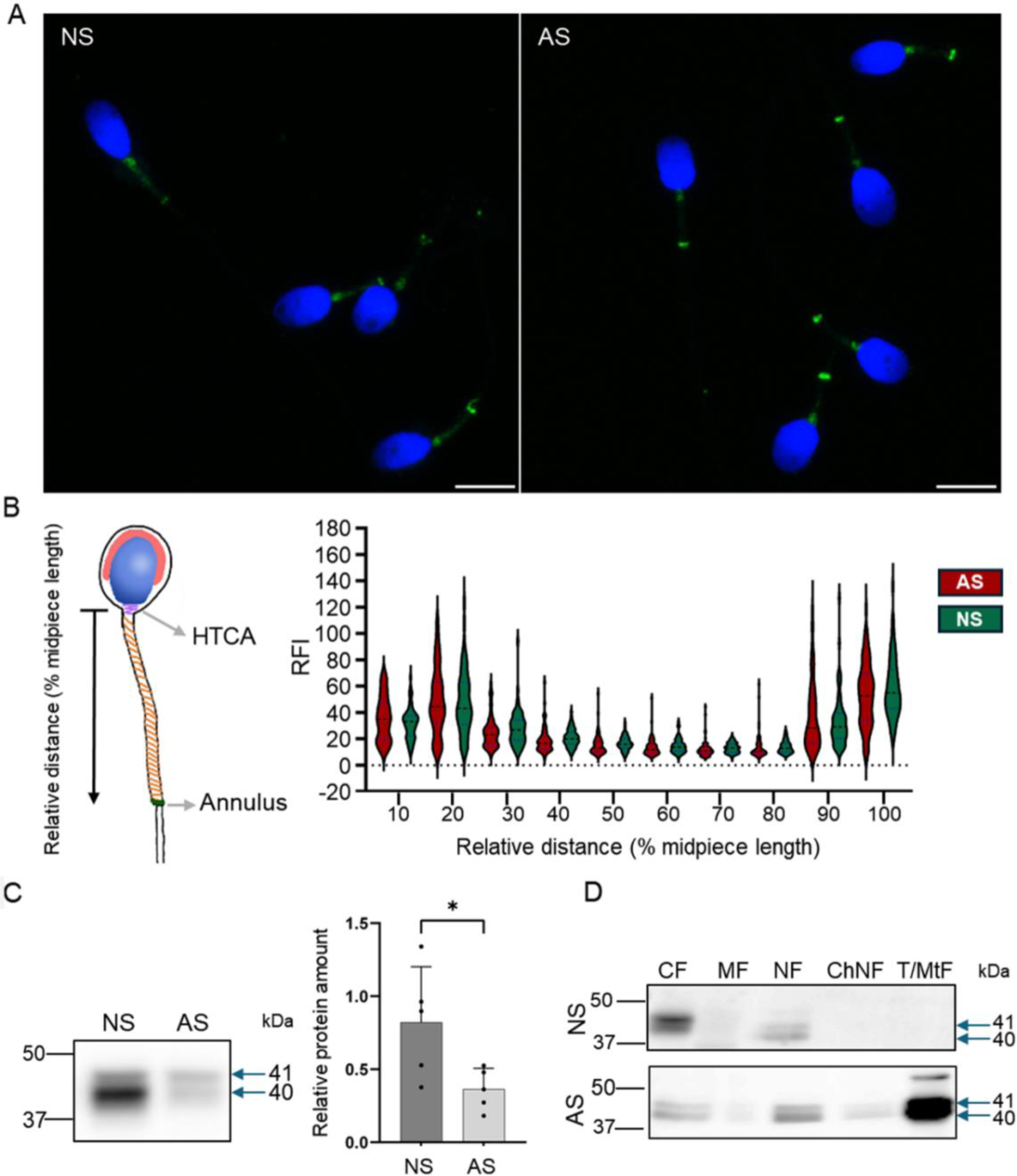
Septin 12 distribution is modified in human asthenozoospermic sperm. **(A)** Comparison of localization of Septin 12 between NS (normozoospermic) and AS (asthenozoospermic) men. Scale bar = 10 µm. **(B)** Graphical visualization of the relative length measurement. The graph shows changes in the relative fluorescence intensity of the Septin 12 signal in the sperm midpiece. Measured relative fluorescent intensity (RF) was compared based on the relative distance (percentage) from the neck. n=3, at least 20 sperm from each patient were measured. T-test, no significant results. **(C)** Western blot shows a decreased abundance of Septin 12 in AS sperm. Blue arrows indicate the molecular size of antibody-detected bands. The graph represents Relative protein amount quantification. n=5, Mann-Whitney test, p=0.0317. **(D)** Subcellular fractionations of human sperm and western blot detection of Septin 12. CF - Cytoplasmic fraction; MF - Membrane fraction; NF - Nuclear fraction; ChNF - Chromatin-bound nuclear fraction. T/MtF -Tubulin/mitochondria protein fraction.

### Septin 12 is retained in asthenozoospermic human sperm

In human sperm, Septin 12 is localized in the nucleus edge (Lin et al., 2011a), likely associated with the nuclear membrane, connecting piece, mitochondrial sheet, and annulus (Lin et al., 2011b). Previous studies reported a decreased level of Septin 12 in men suffering from asthenozoospermia and correlated Septin 12 malfunction with sperm morphology abnormalities (Kuo et al., 2012, Ihara et al., 2005, Sugino et al., 2008, Lhuillier et al., 2009). Notably, patients with asthenozoospermia exhibited differences in the protamine 1 to protamine 2 ratio compared to the control group (Mengual et al., 2003). We used immunofluorescent labeling to visualize Septin 12 localization in human sperm (Fig. 9) and detected it predominantly in connecting piece and annulus in normozoospermic donors and asthenozoospermic patients’ samples. A weak signal was also detected across the whole midpiece of the tail, where mitochondria are located (Fig. 9A). To compare the Septin 12 accumulation within sperm tail of asthenozoospermic and normozoospermic men, relative fluorescent intensities (RFI) were calculated using Zen Blue software, which did not detect change in the localization of Septin 12 in the annulus, or midpiece analyzing a signal between the neck and annulus (Fig. 9B). Subsequent analysis of fluorescent data and normalization to relative length (percentage of the distance) did not again provide a significant change among the patients and healthy donors.

Based on our *Prm2^-/-^* results reporting differences in the Septin 12 isoforms (40 and 41 kDa) compared to WT males, we determined the total Septin 12 protein level using western blot densitometry, normalized to tubulin, in human samples. The analyzed data revealed a significantly (p=0.0317) decreased level of detected Septin 12 isoforms in asthenozoospermic patients (Fig. 9C). Using in-depth analysis of subcellular localization of Septin 12 we isolated subcellular protein fractions and detected a strong signal for Septin 12 (isoforms of 40 and 41 kDa) only in cytoplasmic and nuclear fractions in normozoospermic donor samples (Fig. 9D). In comparison with asthenozoospermic patients’ sperm, Septin 12 was additionally detected in the membrane and nuclear-bound protein fractions apart from the other fraction of normozoospermic donors (Fig. 9D) with very strong signal in fraction defined as tubulin/mitochondria proteins. The retention of Septin 12 in the nuclear-bound fraction of asthenozoospermic men may have a significant value and possibly point to impaired docking of Septin 12 during spermiogenesis, contributing to poor sperm motility.

### Results summary

The schematic illustration (Fig. 10) summarizes the results of this study in connection to current available knowledge relevant to the scope of this work. It describes the connection between Prm2^-/-^ phenotype and Septin 12 and highlights the new possible pathways and missing links. Our results demonstrate distinct localization patterns of Septin 12 individual isoforms in sperm, suggesting that these isoforms could play different roles in sperm organelle biogenesis and function, and the absence of Septin 12 isoforms in the chromatin-bound nuclear protein fraction of *Prm2^-/-^* sperm could be due to Protamine 2 deficiency. Based on co-immunoprecipitation detecting interactions between Septin 12 and Lamin B2/3, we hypothesize that in the absence of Protamine 2, the interplay between Lamin B2/3 and other nuclear lamina-associated proteins might be modified. The differences in the protein abundance pattern of Septin 12 isoforms in caudal sperm may further suggest a role of Septin 12 during the late stages of sperm development and/or during the epididymal maturation. Based on these findings, we propose that Septin 12 and Protamine 2 could potentially directly or indirectly interact within the sperm nucleus.

**Figure 10:**
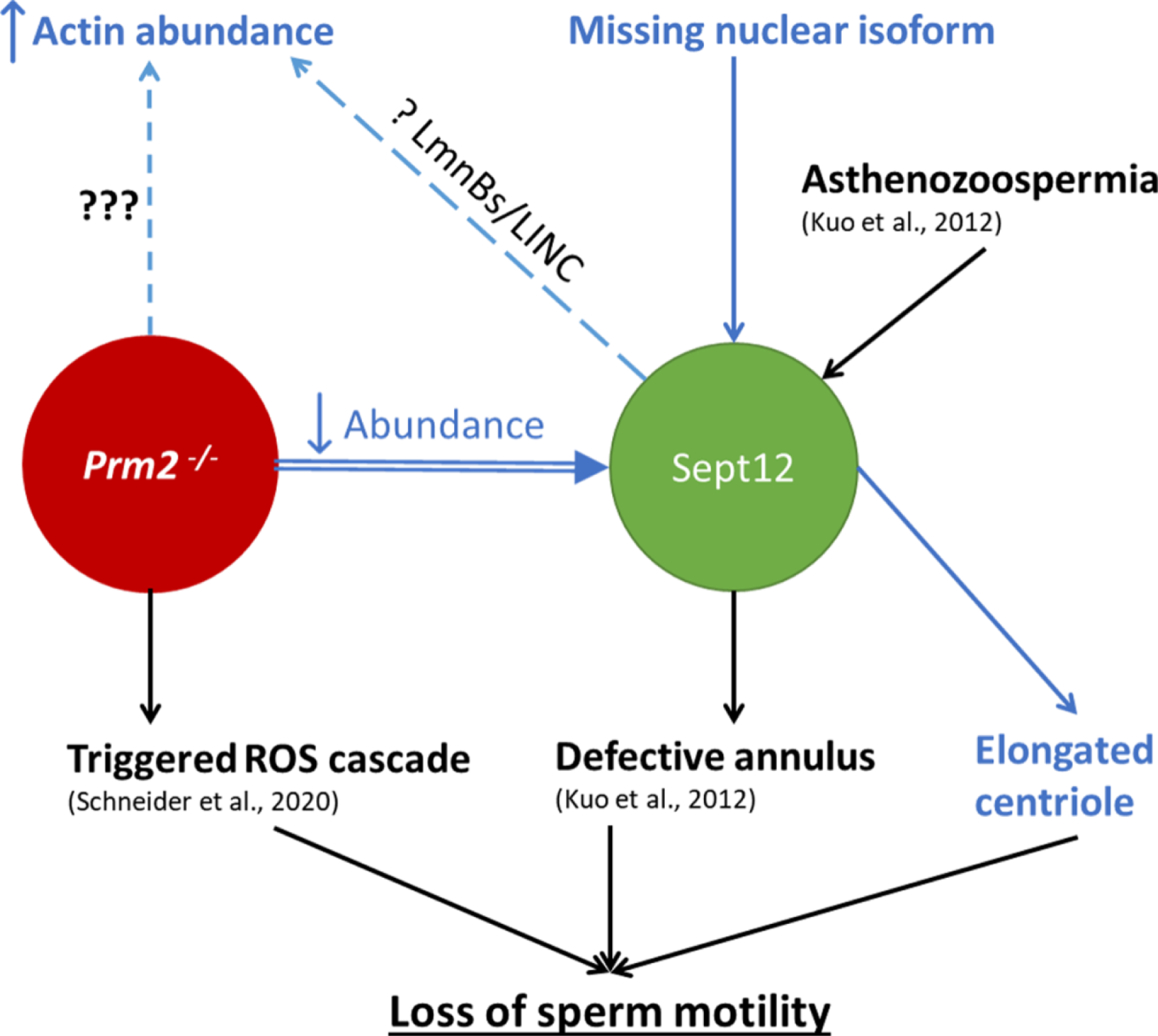
Schematic illustration summarizing the current knowledge regarding the connection between Prm2^-/-^ phenotype and Septin 12 role in mouse sperm. The text in blue represents the results delivered by this study. Arrows up/down represent increased/decreased effect on the described mechanism, dashed lines represent a phenotype without a described cause.represents an unknown cause of the abnormality; The question mark (?) represents a possible mechanism behind the abnormality.

## Discussion

Protamines are known to play a pivotal role in sperm function, its fertilizing ability, and proper early embryogenesis (Arévalo et al., 2022), and Protamine 2 is one of the most abundant sperm proteins in humans and mice (Retief and Dixon, 1993). The establishment of *Prm2^-/-^* mouse strain has unveiled a notable link between Protamine 2 deficiency and male infertility characterized by morphologically impaired and immotile sperm (Schneider et al., 2016), and a follow-up study (Schneider et al., 2020) suggested a possible molecular mechanism behind described sperm dysfunction. Our current study provides new evidence suggesting that Protamine 2 deficiency might significantly impact the cytoskeleton function, especially the testis-specific Septin 12 protein, whose impairment could likely contribute to the development of morphological abnormalities and immotility of *Prm2^-/-^*sperm.

In this study, we employed the DNA FCM analysis and showed differences in DNA hypercondensation during spermiogenesis between WT and *Prm2^-/-^* males and discovered an absence of sub-haploid cells in *Prm2^-/-^* mouse testis compared to WT. Additionally, we investigated the potential impact of a lack of Protamine 2 on histone replacement and retention, including the abundance of a core histone and related epigenetic modifications. While the proteomic approach indicates that histone retention (Arévalo et al., 2022) and DNA methylation may be affected in sperm with aberrant protamine replacement (Carrell et al., 2008), our new approach using FCM analysis did not reveal significant differences in histone H3 retention, or H3K27Me3, H3K36Me3, and H3K9Ac histone pattern between WT and *Prm2^-/-^*sperm. This finding may suggest a dominant role of Protamine 1 in sperm DNA packaging in mice but also a need for further in-depth studies of sperm epigenome and DNA structure to visualize Protamine 1 coils assembly and histone localization in the absence of Protamine 2.

Morphological characterization of *Prm2^-/-^* germ cells did not indicate significant differences in nuclear shape or acrosome biogenesis between *Prm2^-/-^* and WT testicular sperm (Schneider et al., 2020, Schneider et al., 2016). The first detectable morphological changes of *Prm2^-/-^* sperm were described during sperm passage through the *epididymis* (Schneider et al., 2016), which led to the conclusion that morphological aberrations of *Prm2^-/-^* sperm are predominantly acquired during epididymal maturation (Schneider et al., 2020). In accordance with this knowledge, our study confirmed a significant increase in the number of abnormal sperm during sperm epididymal transit in *Prm2^-/-^* mice, and we further described several serious morphological aberrations in caudal *Prm2^-/-^*sperm compared to WT. Interestingly, our examination of testicular sperm acrosomes revealed their abnormal morphological shapes and modified structural appearance, i.e., before the epididymal maturation. These findings suggest a new role of Protamine 2 in spermatid development, which impacts acrosome biogenesis as a critical part of spermiogenesis. At this stage of sperm development, a formation of the acrosome and spermatid head shaping is orchestrated by the cytoskeleton network and represents a crucial process that also involves sperm motility. Identifying abnormal acrosomal shaping in testicular sperm of *Prm2^-/-^* mice, coupled with the observation of complete immotility in *Prm2^-/-^* sperm, points towards a potential impairment of cytoskeletal protein function.

Among other cytoskeletal proteins and their multiple roles, the organisation of actin and tubulin network in sperm is facilitated by testis-specific Septin 12 (Nakos et al., 2022) a member of the Septin cytoskeleton family (Mostowy and Cossart, 2012). In both WT and *Prm2^-/-^*sperm immunofluorescent staining for Septin 12 was located to the neck, annulus with moderate signal over the tail midpiece, and western blot data analysis showed significant differences in Septin 12 isoforms of 40 and 41 kDa corresponding to calculated molecular weights of Septin12 isoforms 3 and X1, which was absent in *Prm2^-/-^* sperm lysate, compared to WT. Subsequent assessment of Septin 12 subcellular localization revealed the presence of these two isoforms in the cytoplasmic fraction of both WT and *Prm2^-/-^* sperm but notably absent from the chromatin-bound nuclear protein fraction of Prm2^-/-^ sperm. These findings suggest potential crosstalk between Septin 12 and Protamine 2 within the sperm nucleus, which aligns well with the reported interaction between Septin 12 and Protamine 2 using a yeast 2-hybrid system (Yeh et al., 2015) and our co-immunoprecipitation results in mouse testes lysate. Consistent with these observations are also our modeling results suggesting interactions between Septin 12 isoforms and Protamine 2. The predicted complexes could be stabilized by potential disulfide bridges involving different Protamine 2 Cysteine residues for each Septin 12 isoform. However, these interactions are still waiting for a more detailed experimental validation. Based on current knowledge, it might be plausible to believe that Septin 12 isoforms are likely to play different roles in sperm function, suggesting regulation of protein localization during later stages of sperm maturation. Septin 12 was previously shown to be essential for correct annulus formation (Lin et al., 2009). Its dysregulation by constitutive phosphorylation leads to immotile sperm with absent annulus. We employed cryo-transmission electron microscopy to study the annulus localization, size, and its distance. Contrary to Shen’s observation, we can see the annulus of *Prm2^-/-^* to be present in the same position as in the WT. Moreover, its area and distance from each other are the same as in WT, even though the immunofluorescent microscopy showed the absence of Septin 12 form *Prm2^-/-^* annulus. These observations might suggest the Septin12 mislocalization during later stages of sperm maturation, potentially as a result of high oxidative *Prm2^-/-^* stress of sperm.

In relevance to the modified status of the cytoskeleton in response to Protamine 2 depletion, a significant increase in the abundance of β-actin was detected in *Prm2^-/-^* sperm lysate. The β-actin and its capping proteins are known to be abnormally regulated in patients with oligozoospermia and asthenozoospermia (Inagaki et al., 2021) and an abnormal regulation of testis-specific actin capping protein CPβ3 was described in subfertile patients (Soda et al., 2017). Moreover, the expression of F-actin-capping protein subunit beta (CAPZB) and Tektin 2 were significantly decreased in patients with low sperm motility, being supported by knowledge that CAPZB causes actin depolymerization meanwhile, tektins provide stability to axonemal microtubules, making these two proteins key factors in sperm motility (Xiong et al., 2018). To add more, the pivotal role of the septin filaments in the anchoring of actin filaments has been reported recently (Martins et al., 2023). Considering the above-reported abnormal acrosomal formation, which is dependent on the cytoskeletal network and the significantly elevated level of β-actin may be a result of miscommunication between cytoskeletal proteins and their associated proteins or from improper sequestration of septin membrane-anchored actin fraction during the spermiogenesis.

Notably, centrioles, which play a pivotal role in regulating sperm tail movement (Avidor-Reiss et al., 2020), exhibit changed morphology under Protamine 2 depletion. This aberration could represent a potential mechanistic link to the observed motility defects and in connection with the previously documented plasma membrane defects resulting in an exceptionally low cytoplasmic Ca^2+^ concentration, may contribute to the observed immotile phenotype (Schneider et al., 2016). This finding is in accordance with results from the yeast 2-hybrid system, which predicted interaction between Septin 12 and PCM1 (Yeh et al., 2015), a component of centriolar satellites (Dammermann and Merdes, 2002). It has been shown that PCM1 and centriolar satellites play a role in regulating the composition of centrioles (Hall et al., 2023) and conversely, PCM1 requires centrosomal localization of CEP135 for efficient localization around centrosomes (Chu and Gruss, 2022). It suggests that the interaction between Septin 12 and PCM1 may regulate the structure and function of sperm centrioles and that the undetected expression of Septin 12 in the connecting piece of *Prm2^-/-^* sperm could indirectly result in their immotility.

Septins, in general, tend to have various interaction partners (Desterke and Gassama-Diagne, 2019). Our results from HEK cells co-immunoprecipitation show the interaction of Septin 12 with Lamin B1, B2, and B3, suggesting its possibly crucial role in spermiogenesis. As described previously, Septin 12 can form a complex with Lamin B1 and SUN4 (a sperm-specific component of LINC complex) and is important for its correct positioning (Lin et al., 2009, Yeh et al., 2015, Yeh et al., 2019). Moreover, (Thoma et al., 2023) it has recently been shown that Lamin B3 interacts with SUN4 *in vivo,* but its localization is independent of SUN4, as shown on the *SUN4^-/-^* mouse model. Based on our analysis, the amount of Lamin B2 and Lamin B3 in testicular cells of *Prm2^-/-^* remains unchanged compared to WT. Similarly, the localization of Lamin B2 and SUN4 in spermatids seems not to be affected by Protamine 2 depletion. Possibly, Septin 12 could be involved in the signal transduction between the nucleus to cytoplasm via the LINC complex, which can be disrupted despite the functional assembly of the LINC components and due to complex multiprotein machinery (revised in (Kmonickova et al., 2020)) further research addressing this topic is needed.

In an attempt to test our findings in relevance to the described clinical observation and human sperm pathologies, we aimed to test Septin 12 protein expression in sperm of asthenozoospermic patients and compare them with normozoospermic donors. Previous studies showed a connection between point mutations of the GTP binding domain of Septin12 with teratozoospermia (Lin et al., 2012), asthenoteratozoospermia, and oligoasthenozoospermia (Kuo et al., 2012). Moreover, the sperm of these patients possessed defective annulus. We discovered that the amount of two isoforms (40 and 41 kDa corresponding to calculated molecular weights of human Septin12 isoform of 2 and X1) of Septin 12 was reduced in asthenozoospermic patients, which is in accordance with previously reported data (Kuo et al., 2012). In contrast to Septin 12 overall reduction, we detected retention of Septin 12 isoforms in subcellular fraction containing specifically tubulin/mitochondrial proteins, and fractions with membrane and chromatin-bound proteins, whereas in normozoospermic donors’ sperm there is none. Humans express less amount of Septin 12 isoforms compared to the expressed in the mouse, so it is likely that their role varies, and a comparison cannot be drawn between human and mouse Septin 12/Protamine 2 related phenotypes. However, the discovery of different localization of Septin 12 isoforms in sperm of asthenozoospermic and normozoospermic men expands our knowledge and contributes to a better understanding of the mechanism behind multiple causes, including idiopathic infertility, behind poor sperm motility in humans.

## Conclusion

In conclusion, our present study provides new insights into sperm physiological abnormalities and immotility caused by Protamine 2 deficiency in mice. Detection of Septin 12 and Protamine 2 interaction in mouse testes, the absence of nuclear Septin 12 isoforms of 40 and 41 kDa in *Prm2^-/-^*sperm as well as the interaction of Septin 12 and Lamin B1/B2/B3 *in vitro* delivers new insight into the role of Protamine 2 in sperm biogenesis. The abundance and localization of Lamin B2 nor Lamin B2/3, as well as localization of SUN4, which were not detected to be influenced by Protamine 2 deletion, suggest a possible new role of Septin 12, which could be involved in the signal transduction between the nucleus to cytoplasm via the LINC complex. This signaling pathway may be disrupted due to Septin 12 modifications despite the functional assembly of the LINC components. This new consequence of Protamine 2 deficiency is also strengthened by the discovery of aberrant acrosome formation during spermiogenesis and results in several cytoskeletal abnormalities that might be caused by aberrant Septin 12 docking.

## Data Availability Statement

Source data files corresponding to individual panels in Figures are deposited in as Source Data Archives: https://zenodo.org/records/11217661.

## Acknowledgment

We acknowledge Michala Krejci for excellent technical assistance, David Liebl and Jiri Miksatko for cryo-TEM data acquisition, Zuzana Cockova for cry-TEM data analysis consultation, and Robert Selvek for statistical data analysis consultation.

This work was supported by the Czech Grant Agency (GC20-20217J) and German Research Foundation (DFG, STE 892/20-1); by institutional support from the Institute of Biotechnology of the Czech Academy of Sciences RVO (86652036); by BIOCEV project (CZ.1.05/1.1.00/02.0109) from the ERDF; European Regional Development Fund, project “Modernization and support of research activities of the national infrastructure for biological and medical imaging Czech-BioImaging” (No. CZ.02.1.01/0.0/0.0/16_013/0001775) Imaging Methods Core Facility at BIOCEV, institution supported by the MEYS CR (Large RI Project LM2018129 Czech-BioImaging) and MEYS CR (LM2023050 Czech-BioImaging); COST Action CA20119 (ANDRONET) supported by European Cooperation in Science and Technology (www.cost.eu). Eunice Kennedy Shriver National Institute of Child Health and Human Development (United States) 1R15HD110863 supported Tomer Avidor-Reiss research.

## Authors contribution

- Conceptualization: OSa, MF, LD, PP, KS, KK
- Data curation: OSa, MF
- Formal Analysis: OSa, VK, OSi, DLC
- Funding acquisition: KS, KK
- Investigation: OSa, MF, VK, MQ, DS, LM, DLC, JC
- Methodology: OSa, MF, VK, JV, MQ, OSi
- Project administration: OSa, KK
- Resources: FL, HS
- Supervision: LD, MF, TAR, SSH, PP, KS, KK
- Validation: VK, JV, LM
- Visualization: OSa, MF, JC
- Writing – original draft: OSa, MF, KK
- Writing – review & editing: TAR, SSH, HS, PP, KS, KK, JC

## Material and methods

### Animals

*Protamine 2* knock-out mouse line (*Prm2^-/-^*) was prepared by Schneider and his colleagues in 2016 (Schneider et al., 2016). Using CRISPR/Cas9 system, 97 bp deletion was introduced into *Prm2* exon 1 between 120-217 bp resulting in premature termination codon occurrence (Fig. S4). This mouse line was registered as C57BL/6J Prm2Δ^97^. Wild-type C57BL/6J males served as a control. Mice were housed in a breeding colony of the Laboratory of Reproduction, IMG animal facilities, Institute of Molecular Genetics of the Czech Academy of Sciences; food and water were supplied ad libitum. The male mice used for all experiments were healthy, 10-14 weeks old, with no sign of stress or discomfort. All animal procedures and experimental protocols were approved by the Animal Welfare Committee of the Czech Academy of Sciences, Animal Ethical protocol code 66866/2015-MZE-17214, 18 December 2015.

### Mouse sperm preparation

Sperm were retrieved from *caput, corpus or cauda epididymis*. The exact sperm source is listed in the corresponding results section. Appropriate parts of the epididymis were dissected, and sperm were released into two 200 μl droplets of M2-fertilizing medium (M7167) under paraffin oil (P14501, P-LAB, Prague, Czech Republic) in a Petri dish and pre-tempered at 37 °C in the 5% CO_2_ atmosphere. After 15 min, medium with released sperm was collected into the Eppendorf tube and centrifuged for 5 min at 300 x g. The supernatant was removed, and the pellet was gently resuspended in 500 µl PBS tempered to 37°C and centrifuged again.

### Human sperm samples

Human ejaculates were obtained from men after 3-4 days of sexual abstinence, at the Centres for Assisted Reproduction (Prague, Czech Republic) with the informed consent of the donors and in accordance with the approval of the Ethical Committee, protocol code BIOCEV 012019, 20 January 2019. Biological materials and experimental protocols were approved by the Ethical Committee of the General University Hospital (Prague, Czech Republic), protocol code 617/17 S-IV.

### Immunofluorescence of mouse and human sperm

For immunofluorescence analysis, the sperm smears were fixed for 10 min with 3.2 % paraformaldehyde and permeabilized with the Intracellular Staining Perm Wash Buffer for 3 x 5 min (421002, BioLegend, USA). Following permeabilization, the samples were blocked for 30 min with SuperBlock blocking buffer (Invitrogen) at room temperature (RT). After the anti-Septin12 primary antibody (H00124404-B01P, Bio-Techne, USA) diluted 1:50 in the Antibody diluent (Zytomed Systems GmbH) was applied and sperm smears were incubated at 4°C overnight. The next day, the slides with sperm smears were washed three times with PBS and then incubated with AlexaFluor 488 goat anti-mouse secondary antibody diluted 1:300 (A11029, Invitrogen, USA) for 60 min and with PNA 568 (Peanut Agglutinin, 6.66 µg/ml, Invitrogen, USA) for 20 min at RT. After, the samples were washed with PBS and incubated with DAPI (62248, Thermo Fischer Scientific, USA) diluted in PBS to the concentration of 2 µg/ml for 5 min at RT. Slices were mounted with AD-Mount-F Mounting Medium (ADVI, Czech Republic) and sealed in coverslips. Images were taken by a laser scanning confocal microscope Carl Zeiss LSM 880 NLO, Zeiss, Germany, Objective Plan-Apochromat 420782-9900(Heide Schatten), magnification 63x, N.A. 1,4.

### Immunohistochemistry of the formalin-fixed paraffin-embedded testes sections

Mouse testes were harvested and fixed in 10% buffered formalin (Sigma-Aldrich, USA) overnight at 4°C. Fixed testes were transferred to 70% ethanol and embedded in paraffin (FFPE blocks). The 5 µm sections were prepared using the Leica microtome system. Testicular FFPE sections were deparaffinized using DIASOLV (DiaPath, Italy) two times for 5 min. After paraffine removal, samples were hydrated in decreasing ethanol series (100%, 90%, 70%) for 5 min in each solution followed by rinsing in distilled water and PBS. Antigen retrieval was performed in a pressure cooker for 15 min in 0.1 M citrate buffer pH 6. After, samples were cooled down to RT for 30 min. After, samples were permeabilized with the Intracellular Staining Perm Wash Buffer for 3 x 5 min (421002, BioLegend), followed by blocking in Superblock solution (Invitrogen, USA) and incubated with antibodies for 1 hour at RT in the humid chamber. Primary antibodies against Lamin B2 (AB_2533107, Thermo Fisher Scientific) and SUN4 (sc-393115, Santa Cruz, USA) were diluted 1:100 and 1:50, respectively, in the Antibody diluent (Zytomed Systems GmbH, Germany) and incubated overnight at 4 °C. The next day, samples were washed in PBS 3 x 5 min at RT with orbital shaking at 100 RPM. Secondary antibodies (Alexa Fluor, Invitrogen) were diluted 1:300 in Antibody diluent and incubated for 1 hour in a humid chamber at RT. Samples were washed twice again. Next, DAPI at a concentration of 2 µg/ml was applied for 5 min to stain the nuclei followed by washing for 5 min in PBS again. Finally, slides were rinsed in distilled water, air-dried, and mounted using the AD-Mount-F Mounting Medium (ADVI, Czech Republic) and sealed with nail polish. Microscopic data were acquired using Carl Zeiss LSM 880 NLO, Zeiss, Germany, Objective Plan-Apochromat 420782-9900(Heide Schatten), magnification 63x, N.A. 1,4 and processed in ImageJ/FIJI and Imaris software.

### 3D visualization of individual seminiferous tubules

Individual seminiferous tubules (iSTs) were processed using the CLARITY procedure (Chung et al., 2013). Briefly, iSTs were fixed in CLARITY gel monomer containing 4% formaldehyde for 2 hours. After gel polymerization occurred, the iSTs were cleared overnight in a clearing buffer (20 mM lithium hydroxide monohydrate and 200 mM SDS, pH 8.5) at 37°C with gentle shaking. The next day, iSTs were stained by DAPI (2µg/ml) and PNA 488 (Thermo Fisher Scientific) (2 µg/ml). Stained iSTs were mounted in PROTOS refractive index matching solution. Z-stacks were acquired using Carl Zeiss LSM 880 NLO confocal microscope, C-Apochromat 421867-9970(Heide Schatten), magnification 40x, N.A. 1,1. For 3D visualization and surface rendering the Imaris software, Version 9.8.2 Package for Cell Biologists was utilized.

### Cryo-Transmission electron microscopy

4 µl of caput sperm suspension were applied to a glow-discharged Quantifoil cryoEM grid (R 2/1, 300mesh) and incubated for 10s in the Leica GP2 Plunge Freezer at room temperature and 80% relative humidity. The grid was then automatically blotted using filter paper for 4s and frozen by immersion into the liquid ethane cooled down to -180°C.

CryoEM data were acquired in a semi-automated mode using SerialEM software (Mastronarde, 2005) on a Jeol JEM 2100Plus transmission electron microscope (TEM) equipped with a LaB6 electron gun and TVIPS XF416 CMOS camera, operated at 200kV. Overview images were taken at magnification x8000 (pixel size 14.04Å) and details were acquired at x20000 (pixel size 5.89Å, electron dose ∼40e/Å2).

### Sperm nuclear proteins isolation and separation

To extract and detect cysteine-containing protamines and histones, we used an optimized protocol from (Shechter et al., 2007, de Yebra et al., 1993). Input for protein extraction was 20 x 10^6^ human sperm for each sample. After sperm washing samples were centrifuged at 12,000 x g at 4°C for 5 min and washed twice in 200 μl of dH2O with Protease inhibitor mix (PIM). The sperm pellet was resuspended in 100 μl of 0.1 M Tris buffer with PIM. After, 100 μl of fresh 6 M Guanidinhydrochloride with 575 mM DTT were added to break disulfide bridges of the nucleo-protamine complex. Next, samples were vortexed and, 200 μl of 255 mM Sodium iodoacetate was added and samples were vortexed again followed by 30 min incubation at RT in the dark. Then, 1 ml of ice-cold 100% EtOH was added, mixed well with samples, and incubated for 1 min at -20 °C. After incubation, samples were centrifuged at 12,000rpm at 4°C for 15 min, and supernatant was removed. The pellet was resuspended in 500 μl of 0.5 M HCl and incubated at 37 °C for 7.5 min (Thermomixer). Incubation was followed by centrifugation at 10,000 x g at 4°C for 10 min. The supernatant was transferred to a new tube (protein low binding tubes, Sarstedt), and 125 μl of 100% trichloracetic acid (TCA) was added, followed by incubation at 4 °C for 10 min. Next, samples were centrifuged again at 14,000 x g at 4°C for 10 min and the protein pellet was washed twice carefully with 1% β-mercaptoethanol in 100% Acetone, centrifugated at 14,000 x g at 4°C for 3 min. The supernatant was removed and Air-dried sperm nuclear protein pellet was resuspended in 20-50 μl of reducing buffer (based on acetic acid containing 0.2% methyl green) and stored at -20 °C.

The protocol was modified according to Shechter et al., Nature Protocols, 2007. Basic nuclear proteins were analyzed using acid–urea polyacrylamide gel electrophoresis. Urea gel containing 2.5 M urea, 0.9 M acetic acid, 15% acrylamide, 0.1% bis-acrylamide, 10% APS, TEMED and H_2_O. After gel polymerization was performed pre-electrophoresis of empty gel in 5% acetic acid (running buffer) for 1-1.5h/150 V in reverse polarity of electrophoretic BioRad system (Bio-rad, USA). The running buffer was discarded, and a fresh one was used for the protein run again with reverse polarity. 10 μl of each sample was loaded and the gel was electrophoresed in 5% acetic acid buffer for 1.5-2h at 150 V.

After the electrophoresis, the gels were fixed in 50% methanol + 10% acetic acid for 15min at RT and washed 3 times in dH_2_O. Consequently, the gels were stained 30-60 min at RT with EzBlue staining reagent (Sigma-Aldrich), following the manufacturer’s instructions, and again washed 3 times in dH_2_O. The stained gels were scanned and the intensity of the bands corresponding to P1 and P2 was quantified with ImageJ/FIJI software. The data obtained from ImageJ/FIJI software were used for calculating the P1/P2 ratio and/or P1 + P2/Histones ratio.

### Testicular cell suspensions preparation and flow cytometry analysis of testicular cells

Testes were harvested and *tunica albuginea* was removed. The rest of the tissue was incubated in RPMI medium with collagenase (35 μg/ml) and DNase (5 μg/ml) at 37 °C, 100 RPM for 20 min. Next, seminiferous tubules were resuspended by gentle pipetting using a wide-end tip. The suspension was filtered through the cell strainer (40 µm, Falcon) a flow-through fraction containing interstitial cells was discarded. Seminiferous tubules were harvested from the strainer and moved into fresh RPMI medium with collagenase (60 μg/ml) and DNase (5 μg/ml) and incubated for 30 min at 37 °C with shaking 150 RPM. After, cells were resuspended by pipetting (20 times) a testicular cell suspension was filtered through the cell strainer (70 µm, Falcon). Flow through fraction containing testicular cells was centrifuged for 10 min, 400 x g at 4 °C. The pellet was resuspended in 1 ml of cold PBS and transferred to a 1.5 ml Eppendorf tube and centrifuged again. The supernatant was removed, and the pellets were gently resuspended in 500 µl of 3.2% PFA in PBS. After 10 min fixation, cells were centrifuged and washed twice in PBS. All centrifugations were performed for 5 min at 400 x g at 4 °C. After the second wash, cell pellets were resuspended in Superblock solution. After 1 hour, cells were counted, and at least 1 x 10^6^ cells were used for subsequent staining. Cells were incubated with primary antibodies (anti-H3 K27me3, ab6002; anti-H3K9ac, ab10812; anti-H3, ab1791; anti-H3K36me3, ab9050) diluted in the Intracellular Staining Perm Wash Buffer 1:100 (BioLegend) overnight at 4 °C. The next day, cells were washed twice with PBS, and goat anti-mouse or anti-rabbit secondary antibody (dilution 1:300) was applied for 1 hour at RT. Finally, cells were stained with DAPI (2 μg/ml) for 5 min and washed twice in PBS. Flow cytometry analyses of the testicular cells were performed using a BD LSRFortessa TM SORP instrument (Becton Dickinson, San Jose, CA, USA). The following lasers and filter parameters were selected, according to which a fluorescent probe was used: FITC blue laser = 488 nm (100 mW) excitation and 530/30 emission filter; PNA 568 = 561 nm (50 mW) laser line excitation and 586/15 nm emission filter; and DAPI = 405 nm (50 mW) laser line excitation and 450/50 nm emission filter. The voltages were set for the optimum resolution, and minimally, 50,000 gated events (based on FSC and SSC) were recorded for each sample. As a standard procedure before each specific parameter evaluation, positive and negative control samples were prepared to ensure the correct setting of the flow cytometer.

### Protein isolation from sperm

To detect Septin 12, the pellet of mouse and human sperm was dissolved in Pellet Extraction Buffer (PEB) with protease inhibitors from the Subcellular Protein Fractionation Kit for Cultured cells (78840, Thermo Fisher Scientific) for 20 min at RT. To detect other proteins, sperm pellets were dissolved in SDS sample buffer (2% SDS, 50 mM Tris-HCl (pH 8), 10 mM EDTA, 10% Glycerol) with Proteinase Inhibitors and incubated on ice for 30 min. Subsequently, the samples were centrifuged at 16,000 x g for 5 min. The supernatant was transferred to a new tube and a 1:1 SDS sample buffer was added to it. The samples were reduced by 5% β-mercaptoethanol and boiled for 5 min at 95 °C. The protein lysate from sperm was used on SDS-Page.

### Protein isolation from mouse testis

Freshly harvested mouse testes were placed in the Precellys tubes with beads (Bertin Technologies, France) and homogenized in corresponding lysis buffer with protease inhibitors by a Precellys tissue homogenizer (5,000 rpm, 10 s, 3 times, 4 °C; Bertin Technologies). Used lysis buffers were: For Septin 12 PEB (from the Subcellular Protein Fractionation Kit for Cultured cells (78840, Thermo Fisher Scientific), form Lamin B2/3 RHB lysis buffer (7M Urea, 2M Thio-urea, 4% CHAPS, 1% Triton X-100, 20mM TRIS) and for other proteins SDS lysis buffer was used. Consequently, the samples were centrifuged at 16,000 × g, for 5 min, at RT, and the supernatant was transferred into a new tube. SDS sample buffer was added to it in a ratio 1:1. The samples were reduced by 5% β-mercaptoethanol and boiled for 5 min at 95 °C. The protein lysate from testis was used on SDS-Page.

### Subcellular fractionation

The Subcellular Protein Fractionation Kit for Cultured cells (78840, Thermo Fisher Scientific) was used for the separation and preparation of cytoplasmic, membrane, nuclear soluble, chromatin-bound, and cytoskeletal protein extracts from mouse and human sperm. The volume of buffers depended on the cell pellet volume, while the volume ratio of used buffers CEB:MEB:NEB:PEB was 200:200:100:100. As a first reagent was added cytoplasmic extraction buffer (CEB), incubated with the sperm for 10 min at 4°C with gentle mixing, followed by centrifugation 500 x g for 5 min at 4°C. The supernatant with cytoplasmic proteins was transferred to a new tube and the membrane extraction buffer (MEB) was added to the pellet. After the 10 min incubation on ice, the samples were centrifuged 3,000 x g for 5 min at 4 °C and the supernatant with membrane proteins was transferred to a new tube. Nuclear extraction buffer (NEB) was added to the pellet, vortexed on higher settings, and incubated with the sample for 30 min on ice. The sample was centrifuged at 5,000 x g for 5 min at 4 °C and the supernatant with nuclear soluble proteins was transferred to a new tube. The NEB with 100 mM CaCl_2_ and 300 units of Micrococcal Nuclease was added to the samples. The samples were incubated for 15 min at RT, followed by centrifugation at 16,000 x g for 5 min at RT. The supernatant with chromatin-bound proteins was transferred to a new tube and PEB was added to the rest of the pellet (Tubulin/mitochondria protein fraction). The sample was incubated for 10 min at RT and centrifuged at 16,000 x g for 5 min. The supernatant was transferred to a new tube and the pellet was discarded. SDS samples buffer was added to the all tubes with supernatant with different subcellular fractions in a ratio 1:1. The samples were reduced by 5% β-mercaptoethanol and boiled for 5 min at 95 °C and the protein fractions were used on SDS-Page.

### SDS Electrophoresis (SDS-Page) and Western Blot Analysis

The protein extracts were separated by 10% SDS-PAGE and the proteins were transferred on the low fluorescence PVDF membrane (Bio-rad). The molecular weight of the proteins was assigned by Precision Plus Protein Dual Color Standards (Bio-Rad). The membranes were blocked in 5% Blotto, non-fat dry milk (Santa Cruz Biotechnology) diluted in PBS and incubated with rabbit polyclonal Septin-12 Antibody (NBP1-91640, Novus Biologicals) diluted 1:250 in PBS, at 4 °C overnight. Other antibodies used in this article were α-tubulin (A11126, Thermo Fisher Scientific 1:600), α-tubulin acK40 (T7451 Sigma-Aldrich, 1:600), β-actin (ab 6276, Abcam, UK, 1:600), β-tubulin (T8355, Sigma-Aldrich, 1:600) and LmnB2/3 (ab151735, Abcam, 1:300). After incubation, the membranes were washed in PBS and incubated with a secondary antibody, goat anti-rabbit IgG or goat anti-mouse IgG conjugated to horseradish peroxidase (Bio-Rad), diluted 1:3,000 in PBS for 1 h at RT, and consequently washed in PBS. The membranes were developed with SuperSignal™ Chemiluminescent Substrate (Thermo Scientific) and images captured using the Azure c600 imaging system (Azure Biosystems, Inc., CA, USA). As a negative control a membrane without a primary antibody was used. The membranes were stained with Coomassie brilliant blue (7% acetic acid, 50% ethanol, dH2O, 0.1% Coomassie brilliant blue) for 5 min at RT and washed in a destaining solution (35% ethanol, 10 % acetic acid, dH2O) to determine the total protein profile.

### Transfection and immunofluorescent staining of HEK293T/17 cells

HEK293T/17 cells were co-transfected transiently with mouse Septin 12 and Lamin B1 or Lamin B2 or Lamin B3. Briefly, 1.1 x 10^6^ cells were seeded in 60 mm cultivation dish containing a coverslip and incubated overnight. The following day, cells with 70-80% confluency were transfected with Lipofectamine 3000 Transfection Reagent (L3000001, Invitrogen) kit using 3 µg of mouse Septin 12-GFP plasmid (pEGFP-C1, Addgene, USA) and 3 µg of mouse Lamin B1 or Lamin B2 or Lamin B3 plasmids conjugated with myc-tag (pCS2-MT, Addgene). Cells were incubated for 48h in a DMEM cultivation medium (Gibco, USA) supplemented with 10% FBS (Gibco, USA) followed by immunofluorescent staining. Briefly, cells on the coverslips were fixed with Paraformaldehyde 3.2% (Scientific) in PBS for 10 min, washed twice with PBS followed by permeabilizing with 0.1% Triton X-100 (Serva, Germany) for 5 min. Super Block (Thermo Fisher Scientific) was used for 1h blocking and cells were incubated with anti-myc tag primary antibody (MA1-21316, Invitrogen) diluted 1:500 for 2h in RT. After 3 times washing, the slides were incubated with donkey anti-mouse IgG secondary antibody conjugated with Alexa Fluor 568 (A10037, Invitrogen) diluted 1:300 for 1h at RT. Subsequently, they were washed with PBS, incubated with DAPI (2 µg/ml) for 5 min, washed in PBS again, dried, and mounted in AD-Mount-F Mounting Medium. Septin 12 was conjugated with a GFP-tag and was not stained. Apochromat 420782-9900 (Heide Schatten), magnification 63x, N.A. 1,4 and processed in ImageJ/FIJI and Imaris software.

### Co-immunoprecipitation (Co-IP)

Co-transfected cells with Septin 12 and Lamin B1 or Lamin B2 or Lamin B3 were incubated for 48h followed by cell lysis with 1% CHAPS (Sigma-Aldrich) in 30 mM Tris-HCl (pH 7.5). Subsequently, Co-IP was performed on cell lysate supernatants by Dynabeads Protein G Immunoprecipitation kit (10007D, Invitrogen). Briefly, magnetic beads were conjugated with anti-GFP antibody (Ab290, Abcam) (against Septin 12) and incubated with cell lysate supernatants overnight at 4°C on a rotator. The samples were washed 3 times and precipitated proteins were eluted in the kit’s elution buffer and incubated at 70°C for 5 min. Rabbit IgG Isotype control antibody (31235, Invitrogen) was used for the isotype control group, and all the procedures were also performed on non-transfected cell lysate supernatant as the control. Then western blot was performed to detect Lamin B1/B2/B3 in the eluted protein complexes as described above. Lamin B1/B2/B3 was detected with anti-myc tag primary antibody (MA1-21316, Invitrogen) diluted 1:500. Freshly harvested mouse testes were placed in the Precellys tubes with beads (Bertin Technologies, France) and homogenized in the 50 mM HEPES buffer with 3 mM MgCl_2_, 500 mM KCl, 20% Glycerol, 1% NP40 (Roche, Switzerland) with protease inhibitors by a Precellys tissue homogenizer (5,000 rpm, 10 s, 3 times, 4 °C; Bertin Technologies, france). The obtained protein lysate was subsequently used for Co-IP of the Prm2-Septin12 protein complex by Dynabeads Protein G Immunoprecipitation kit (10007D, Invitrogen, USA). Magnetic beads were conjugated with anti-protamine 2 antibody (Mab-Hup2B, Labome, USA) for 30 min at RT. As a control, magnetic beads conjugated with a mouse IgG isotype control (31903, Invitrogen) were used. Subsequently, the beads were washed, and lysate obtained from the isolation of mouse testes was added to them. Samples were incubated overnight at 4°C with constant agitation. The next day, the supernatant was removed, the beads with protein complex were 3 times washed and the precipitated proteins were eluted from beads by elution buffer with SDS sample buffer and 5% β-mercaptoethanol was added to the beads and incubated at 70°C for 5 min. Subsequently, SDS-Page with Western Blot Analysis was performed, followed by the detection of Septin 12 Antibody (NBP1-91640, Novus Biologicals, USA).

### Centriole analysis

To analyze signal intensity, length, and width of the centrioles, we used CEP135 immunofluorescent staining using a rabbit polyclonal antibody raised against the first 233 amino acids of CEP135 (24428-1-AP, Thermo Fisher Scientific). Sperm samples were visualized using a Leica SP8 confocal microscope in BrightR mode using an HC PL APO CS2 63x/1.40 OIL lens, 100% gain, 1024 × 1024 pixels (62 μm x 62 μm) format, 3x zoom factor, line averaging of 3, and frame accumulation of 2. Three sequences were used to collect the fluorescence signals. DNA signal was detected using a 410 nm laser set to detect photons between 425-478 nm and was color-coded to blue. For phase-like images, the fluoro-turret was set to Scan-PH, and PMT Trans was set to ON with a gain of 300 in greyscale. CEP135 signal was detected using an anti-rabbit secondary conjugated with ALEXA 488 and activated with a 488 nm laser. The absorption spectrum was set to 505-550 nm and was color-coded to green. The Tubulin signal was detected using an anti-sheep secondary conjugated with ALEXA 555 and activated with a 561 nm laser. We set the absorption spectrum to 560-625 nm, and it was color-coded to red. We collected 10-15 Z-sections of 0.3 μm thickness from the bottom to the top of the sperm. The CEP135 intensity was quantified using a Leica LAS-X algorithm. Centriole length and width were determined using the line-profile tool in Leica LAS-X. A line 3 μm long and 0.5 μm wide was drawn through the CEP135 labeling, and a corresponding graph was calculated for each pixel value along the line. The distance from two points at half the distance of the graph’s peak was measured to determine the length and width of CEP135.

### RNA isolation, reverse transcription, qPCR

Total RNA was isolated from testicular tissue samples using TRI Reagent®. RNA extracts (2 μg) were treated with DNase I (1 U/µl, Fermentas, USA) in the presence of DNase I buffer 10 × (Thermo Fisher Scientific,) with MgCl2 for 30 min at 37 °C, and EDTA (Fermentas) was added for 10 min at 65 °C. The reverse transcription reaction contained 5 × reaction Buffer (Fermentas), RNaseOUT Recombinant Ribonuclease Inhibitor (40 U/1 µL, Thermo Fisher Scientific), Universal RNA Spike II (0.005 ng/µL, TATAA biocenter, Göteborg, Sweden), 10 mM dNTP Mix (Thermo Fisher Scientific), oligo(dt)18 (Thermo Fisher Scientific) mixed 1:1 with Random primers (Thermo Scientific) and M-MuLV RevertAid transcriptase (200 U/µL, Fermentas), and run to generate cDNA. The RT^-^ negative control was prepared in the same conditions but with RNase/DNase-free water. For qRT-PCR, 10 ng/µl cDNA was used. Two times Maxima SYBR Green qPCR Master Mix (Thermo Fisher Scientific), reverse and forward primer and nuclease-free water were used, and all reactions were performed in duplets in a PCR cycler (CFX 96-qPCR cycler / CFX 384-qPCR, Bio-Rad). The RT^-^ negative control for cDNA synthesis, no template control, and spike control were also analyzed. Ribosomal protein S2 (*Rps2*) was used as a reference gene.

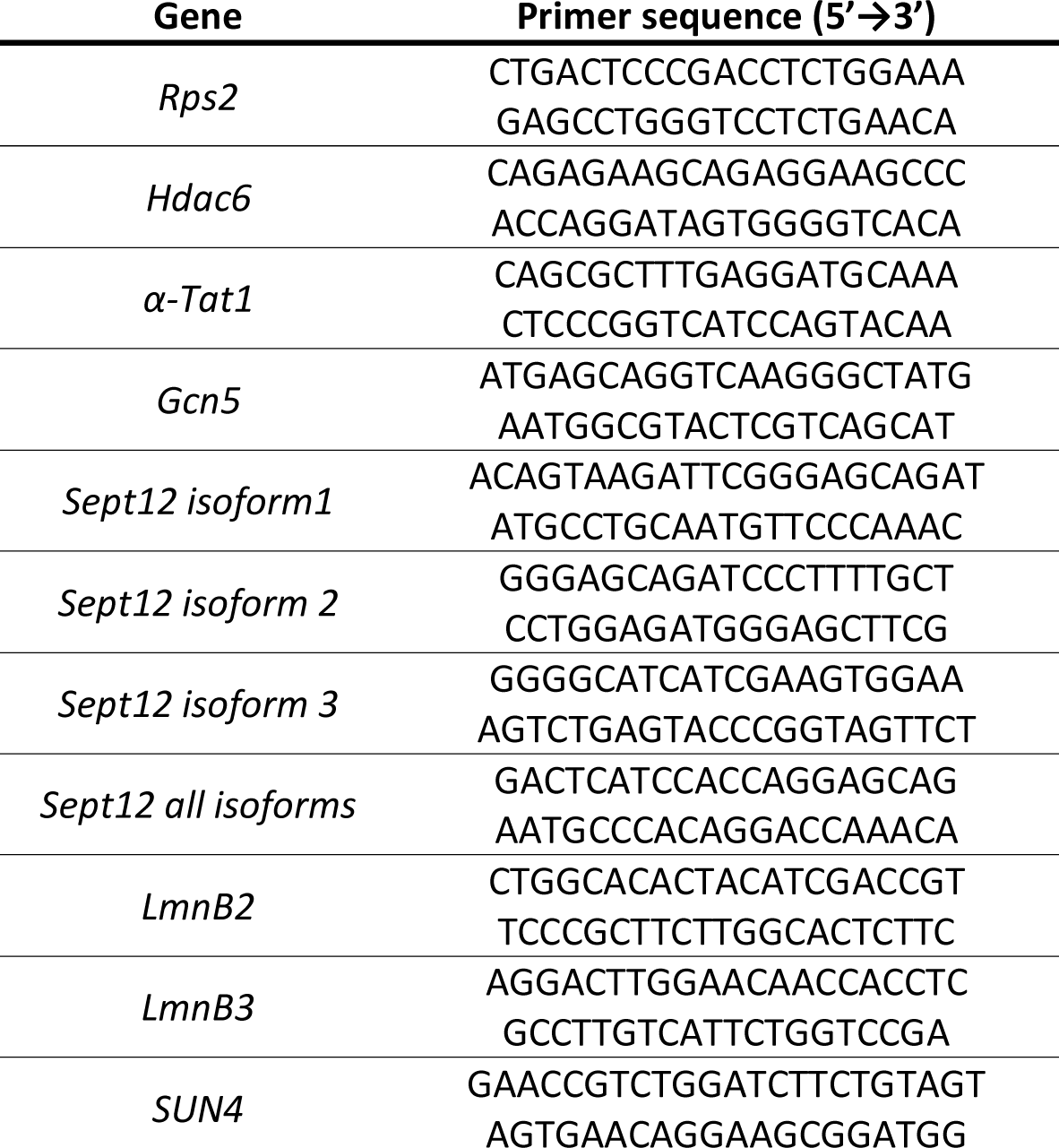

### Statistical and data analysis

The intensity of Septin 12 signal among the tail was analyzed in ZEN Blue software. The line in the width of 5 pixels was drawn for each sperm from the head (0 % of the distance) to the end of the annulus (100 % of the distance). Intensities were normalized on the relative distance from the head (0 %). Measured values were subsequently analyzed using Microsoft Excel. Data from flow cytometry were gated and analyzed using Flow Jo software. The gating strategy is shown in Fig. S1. 3D microscopic data were processed using Imaris (version 9.8.2 Package for Cell Biologists) software. Cryo-TEM data were analyzed in FIJI software using Labkit, MorpholibJ and 3DSuite plugins. Annulus was manually detected, and its area was marked using “masks” function. The distance of annulus centroids (geometric center of a structure) and the area of these structures were. Graphs and statistical analysis were done using GraphPad software (version 10.2.0) using Mann-Whitney or T-test.

### Molecular Modelling

The prediction of potential complexes of Protamine 2 with Septin 12 was performed using a local installation of the AlphaFold version 2.3.2 {10.1038/s41586-021-03819-2}. Three complexes were prepared using AlphaFold multimer with protein sequences of mouse Protamine 2 (NP_032959.1) with Septin 12 isoform 1 (NP_081945.1), mouse Protamine 2 (NP_032959.1) with Septin 12 isoform 3 (NP_001360874.1), and human Protamine 2 (NP_002753.2) with Septin 12 isoform 2 (NP_653206.2), respectively. The modelling results were visualized with the open-source PyMOL version 2.6.0 (The PyMOL Molecular Graphics System, Schrödinger, LLC). The pLDDT scores were transformed in PyMOL to 100-pLDDT for the visualization using the standard B-factor color scale with cold colors corresponding to high confidence regions (see Fig. 8).

## References

Ammer, H., Henschen, A. & Lee, C. H. 1986. Isolation and amino-acid sequence analysis of human sperm protamines P1 and P2. Occurrence of two forms of protamine P2. Biol Chem Hoppe Seyler, 367, 515–22.

Arévalo, L., Merges, G. E., Schneider, S., Oben, F. E., Neumann, I. S. & Schorle, H. 2022. Loss of the cleaved-protamine 2 domain leads to incomplete histone-to-protamine exchange and infertility in mice. PLos Genet, 18, e1010272.

Avidor-Reiss, T., Carr, A. & Fishman, E. L. 2020. The sperm centrioles. Mol Cell Endocrinol, 518, 110987.

Balhorn, R. 1982. A model for the structure of chromatin in mammalian sperm. J Cell Biol, 93, 298–305.

Brunner, A. M., Nanni, P. & Mansuy, I. M. 2014. Epigenetic marking of sperm by post-translational modification of histones and protamines. Epigenetics & Chromatin, 7, 2.

Calvi, A., Wong, A. S., Wright, G., Wong, E. S., Loo, T. H., Stewart, C. L. & Burke, B. 2015. SUN4 is essential for nuclear remodeling during mammalian spermiogenesis. Dev Biol, 407, 321–30.

Carrell, D. T., Emery, B. R. & Hammoud, S. 2008. The aetiology of sperm protamine abnormalities and their potential impact on the sperm epigenome. International Journal of Andrology, 31, 537–545.

Chu, Z. & Gruss, O. J. 2022. Mitotic Maturation Compensates for Premature Centrosome Splitting and Pcm Loss in Human cep135 Knockout Cells. Cells, 11.

Chung, K., Wallace, J., Kim, S. Y., Kalyanasundaram, S., Andalman, A. S., Davidson, T. J., Mirzabekov, J. J., Zalocusky, K. A., Mattis, J., Denisin, A. K., Pak, S., Bernstein, H., Ramakrishnan, C., Grosenick, L., Gradinaru, V. & Deisseroth, K. 2013. Structural and molecular interrogation of intact biological systems. Nature, 497, 332–7.

Coelingh, J. P., Rozijn, T. H. & Monfoort, C. H. 1969. Isolation and partial characterization of a basic protein from bovine sperm heads. Biochim Biophys Acta, 188, 353–6.

Cooper, T. G. 2005. Cytoplasmic droplets: the good, the bad or just confusing? Human Reproduction, 20, 9–11.

Corzett, M., Mazrimas, J. & Balhorn, R. 2002. Protamine 1: protamine 2 stoichiometry in the sperm of eutherian mammals. Mol Reprod Dev, 61, 519–27.

Dammermann, A. & Merdes, A. 2002. Assembly of centrosomal proteins and microtubule organization depends on Pcm-1. J Cell Biol, 159, 255–66.

De Yebra, L., Ballescà, J. L., Vanrell, J. A., Bassas, L. & Oliva, R. 1993. Complete selective absence of protamine P2 in humans. J Biol Chem, 268, 10553–7.

Desterke, C. & Gassama-Diagne, A. 2019. Protein-protein interaction analysis highlights the role of septins in membrane enclosed lumen and mrna processing. Advances in Biological Regulation, 73, 100635.

Dunleavy, J. E. M., O’bryan, M. K., Stanton, P. G. & O’donnell, L. 2019. The cytoskeleton in spermatogenesis. Reproduction, 157, R53–R72.

Erkek, S., Hisano, M., Liang, C.-Y., Gill, M., Murr, R., Dieker, J., Schübeler, D., Vlag, J. V. D., Stadler, M. B. & Peters, A. H. 2013. Molecular determinants of nucleosome retention at Cpg-rich sequences in mouse spermatozoa. Nature structural & molecular biology, 20, 868–875.

Frolikova, M., Sebkova, N., Ded, L. & Dvorakova-Hortova, K. 2016. Characterization of CD46 and β1 integrin dynamics during sperm acrosome reaction. Sci Rep, 6, 33714.

Furukawa, K. & Hotta, Y. 1993. cdna cloning of a germ cell specific lamin B3 from mouse spermatocytes and analysis of its function by ectopic expression in somatic cells. Embo j, 12, 97–106.

Gatewood, J. M., Cook, G. R., Balhorn, R., Bradbury, E. M. & Schmid, C. W. 1987. Sequence-specific packaging of Dna in human sperm chromatin. Science, 236, 962–4.

Hall, E. A., Kumar, D., Prosser, S. L., Yeyati, P. L., Herranz-Pérez, V., García-Verdugo, J. M., Rose, L., Mckie, L., Dodd, D. O., Tennant, P. A., Megaw, R., Murphy, L. C., Ferreira, M. F., Grimes, G., Williams, L., Quidwai, T., Pelletier, L., Reiter, J. F. & Mill, P. 2023. Centriolar satellites expedite mother centriole remodeling to promote ciliogenesis. eLife, 12, e79299.

Hammoud, S. S., Nix, D. A., Zhang, H., Purwar, J., Carrell, D. T. & Cairns, B. R. 2009. Distinctive chromatin in human sperm packages genes for embryo development. Nature, 460, 473–478.

Heide Schatten, P., Vanesa Y. Rawe, PHD2-3, Qing-Yuan Sun, PHD4 2011. The sperm centrosome: its role and significance in nature and human assisted reproduction. J Reprod Stem Cell Biotechno, 2, 121–127.

Hisano, M., Erkek, S., Dessus-Babus, S., Ramos, L., Stadler, M. B. & Peters, A. H. 2013. Genome-wide chromatin analysis in mature mouse and human spermatozoa. Nature protocols, 8, 2449–2470.

Ihara, M., Kinoshita, A., Yamada, S., Tanaka, H., Tanigaki, A., Kitano, A., Goto, M., Okubo, K., Nishiyama, H., Ogawa, O., Takahashi, C., Itohara, S., Nishimune, Y., Noda, M. & Kinoshita, M. 2005. Cortical organization by the septin cytoskeleton is essential for structural and mechanical integrity of mammalian spermatozoa. Dev Cell, 8, 343–52.

Inagaki, Y., Fukuhara, S., Kuribayashi, S., Okada, K., Sekii, Y., Takezawa, K., Kiuchi, H., Soda, T., Miyagawa, Y., Okamoto, Y., Tanaka, H. & Nonomura, N. 2021. The expression of human testis-specific actin capping protein predicts in vitro fertilization outcomes: A novel biomarker of sperm function for assisted reproductive technology. Reprod Med Biol, 20, 537–542.

Khanal, S., Jaiswal, A., Chowdanayaka, R., Puente, N., Turner, K., Assefa, K. Y., Nawras, M., Back, E. D., Royfman, A., Burkett, J. P., Cheong, S. H., Fisher, H. S., Sindhwani, P., Gray, J., Ramachandra, N. B. & Avidor-Reiss, T. 2024. The evolution of centriole degradation in mouse sperm. Nature Communications, 15, 117.

Kierszenbaum, A. L., Rivkin, E. & Tres, L. L. 2003. Acroplaxome, an F-Actin–Keratin-containing Plate, Anchors the Acrosome to the Nucleus during Shaping of the Spermatid Head. Molecular Biology of the Cell, 14, 4628–4640.

Kierszenbaum, A. L., Tres, L. L., Rivkin, E., Kang-Decker, N. & Van Deursen, J. M. A. 2004. The Acroplaxome Is the Docking Site of Golgi-Derived Myosin Va/Rab27a/b-Containing Proacrosomal Vesicles in Wild-Type and Hrb Mutant Mouse Spermatids1. Biology of Reproduction, 70, 1400–1410.

Kmonickova, V., Frolikova, M., Steger, K. & Komrskova, K. 2020. The Role of the Linc Complex in Sperm Development and Function. Int J Mol Sci, 21.

Kuo, Y.-C., Lin, Y.-H., Chen, H.-I., Wang, Y.-Y., Chiou, Y.-W., Lin, H.-H., Pan, H.-A., Wu, C.-M., Su, S.-M., Hsu, C.-C. & Kuo, P.-L. 2012. SEPT12 mutations cause male infertility with defective sperm annulus. Human Mutation, 33, 710–719.

Lhuillier, P., Rode, B., Escalier, D., Lorès, P., Dirami, T., Bienvenu, T., Gacon, G., Dulioust, E. & Touré, A. 2009. Absence of annulus in human asthenozoospermia: case report. Hum Reprod, 24, 1296–303.

Lin, Y. H., Chou, C. K., Hung, Y. C., Yu, I. S., Pan, H. A., Lin, S. W. & Kuo, P. L. 2011a. SEPT12 deficiency causes sperm nucleus damage and developmental arrest of preimplantation embryos. Fertil Steril, 95, 363–5.

Lin, Y. H., Kuo, Y. C., Chiang, H. S. & Kuo, P. L. 2011b. The role of the septin family in spermiogenesis. Spermatogenesis, 1, 298–302.

Lin, Y. H., Lin, Y. M., Wang, Y. Y., Yu, I. S., Lin, Y. W., Wang, Y. H., Wu, C. M., Pan, H. A., Chao, S. C., Yen, P. H., Lin, S. W. & Kuo, P. L. 2009. The expression level of septin12 is critical for spermiogenesis. Am J Pathol, 174, 1857–68.

Lin, Y. H., Wang, Y. Y., Chen, H. I., Kuo, Y. C., Chiou, Y. W., Lin, H. H., Wu, C. M., Hsu, C. C., Chiang, H. S. & Kuo, P. L. 2012. SEPTIN12 genetic variants confer susceptibility to teratozoospermia. PLos One, 7, e34011.

Maier, W. M., Nussbaum, G., Domenjoud, L., Klemm, U. & Engel, W. 1990. The lack of protamine 2 (P2) in boar and bull spermatozoa is due to mutations within the P2 gene. Nucleic Acids Res, 18, 1249–54.

Martins, C. S., Taveneau, C., Castro-Linares, G., Baibakov, M., Buzhinsky, N., Eroles, M., Milanović, V., Omi, S., Pedelacq, J. D., Iv, F., Bouillard, L., Llewellyn, A., Gomes, M., Belhabib, M., Kuzmić, M., Verdier-Pinard, P., Lee, S., Badache, A., Kumar, S., Chandre, C., Brasselet, S., Rico, F., Rossier, O., Koenderink, G. H., Wenger, J., Cabantous, S. & Mavrakis, M. 2023. Human septins organize as octamer-based filaments and mediate actin-membrane anchoring in cells. J Cell Biol, 222.

Mastronarde, D. N. 2005. Automated electron microscope tomography using robust prediction of specimen movements. Journal of structural biology, 152, 36–51.

Mengual, L., Ballescá, J. L., Ascaso, C. & Oliva, R. 2003. Marked differences in protamine content and P1/P2 ratios in sperm cells from percoll fractions between patients and controls. J Androl, 24, 438–47.

Mostowy, S. & Cossart, P. 2012. Septins: the fourth component of the cytoskeleton. Nature Reviews Molecular Cell Biology, 13, 183–194.

Mukherjee, A., Saurabh, S., Olive, E., Jang, Y. H. & Lansac, Y. 2021. Protamine binding site on Dna: molecular dynamics simulations and free energy calculations with full atomistic details. The Journal of Physical Chemistry B, 125, 3032–3044.

Nakos, K., Alam, M. N. A., Radler, M. R., Kesisova, I. A., Yang, C., Okletey, J., Tomasso, M. R., Padrick, S. B., Svitkina, T. M. & Spiliotis, E. T. 2022. Septins mediate a microtubule–actin crosstalk that enables actin growth on microtubules. Proceedings of the National Academy of Sciences, 119, e2202803119.

Pereira, C. D., Serrano, J. B., Martins, F., Da Cruz, E. S. O. A. B. & Rebelo, S. 2019. Nuclear envelope dynamics during mammalian spermatogenesis: new insights on male fertility. Biol Rev Camb Philos Soc, 94, 1195–1219.

Retief, J. D. & Dixon, G. H. 1993. Evolution of pro-protamine P2 genes in primates. Eur J Biochem, 214, 609–15.

Rodríguez-Casuriaga, R. & Geisinger, A. 2021. Contributions of Flow Cytometry to the Molecular Study of Spermatogenesis in Mammals. International Journal of Molecular Sciences, 22, 1151.

Rogenhofer, N., Dansranjavin, T., Schorsch, M., Spiess, A., Wang, H., Von Schönfeldt, V., Cappallo-Obermann, H., Baukloh, V., Yang, H., Paradowska, A., Chen, B., Thaler, C. J., Weidner, W., Schuppe, H. C. & Steger, K. 2013. The sperm protamine mrna ratio as a clinical parameter to estimate the fertilizing potential of men taking part in an Art programme. Hum Reprod, 28, 969–78.

Rogenhofer, N., Ott, J., Pilatz, A., Wolf, J., Thaler, C. J., Windischbauer, L., Schagdarsurengin, U., Steger, K. & Von Schönfeldt, V. 2017. Unexplained recurrent miscarriages are associated with an aberrant sperm protamine mrna content. Human Reproduction, 32, 1574–1582.

Schagdarsurengin, U., Paradowska, A. & Steger, K. 2012. Analysing the sperm epigenome: roles in early embryogenesis and assisted reproduction. Nature Reviews Urology, 9, 609–619.

Schagdarsurengin, U. & Steger, K. 2016. Epigenetics in male reproduction: effect of paternal diet on sperm quality and offspring health. Nat Rev Urol, 13, 584–95.

Schneider, S., Balbach, M., Jan, F. J., Fietz, D., Nettersheim, D., Jostes, S., Schmidt, R., Kressin, M., Bergmann, M., Wachten, D., Steger, K. & Schorle, H. 2016. Re-visiting the Protamine-2 locus: deletion, but not haploinsufficiency, renders male mice infertile. Sci Rep, 6, 36764.

Schneider, S., Shakeri, F., Trötschel, C., Arévalo, L., Kruse, A., Buness, A., Poetsch, A., Steger, K. & Schorle, H. 2020. Protamine-2 Deficiency Initiates a Reactive Oxygen Species (Ros)-Mediated Destruction Cascade during Epididymal Sperm Maturation in Mice. Cells, 9.

Schütz, W., Alsheimer, M., Ollinger, R. & Benavente, R. 2005a. Nuclear envelope remodeling during mouse spermiogenesis: postmeiotic expression and redistribution of germline lamin B3. Exp Cell Res, 307, 285–91.

Schütz, W., Benavente, R. & Alsheimer, M. 2005b. Dynamic properties of germ line-specific lamin B3: the role of the shortened rod domain. Eur J Cell Biol, 84, 649–62.

Shechter, D., Dormann, H. L., Allis, C. D. & Hake, S. B. 2007. Extraction, purification and analysis of histones. Nat Protoc, 2, 1445–57.

Soda, T., Miyagawa, Y., Ueda, N., Takezawa, K., Okuda, H., Fukuhara, S., Fujita, K., Kiuchi, H., Uemura, M., Okamoto, Y., Tsujimura, A., Tanaka, H. & Nonomura, N. 2017. Systematic characterization of human testis-specific actin capping protein β3 as a possible biomarker for male infertility. Human Reproduction, 32, 514–522.

Sugino, Y., Ichioka, K., Soda, T., Ihara, M., Kinoshita, M., Ogawa, O. & Nishiyama, H. 2008. Septins as diagnostic markers for a subset of human asthenozoospermia. J Urol, 180, 2706–9.

Thoma, H., Grünewald, L., Braune, S., Pasch, E. & Alsheimer, M. 2023. SUN4 is a spermatid type Ii inner nuclear membrane protein that forms heteromeric assemblies with SUN3 and interacts with lamin B3. J Cell Sci, 136.

Wang, T., Gao, H., Li, W. & Liu, C. 2019. Essential Role of Histone Replacement and Modifications in Male Fertility. Front Genet, 10, 962.

Ward, W. S. & Coffey, D. S. 1991. Dna packaging and organization in mammalian spermatozoa: comparison with somatic cells. Biol Reprod, 44, 569–74.

Xiong, Z., Zhang, H., Huang, B., Liu, Q., Wang, Y., Shi, D. & Li, X. 2018. Expression pattern of prohibitin, capping actin protein of muscle Z-line beta subunit and tektin-2 gene in Murrah buffalo sperm and its relationship with sperm motility. Asian-Australas J Anim Sci, 31, 1729–1737.

Yeh, C. H., Kuo, P. L., Wang, Y. Y., Wu, Y. Y., Chen, M. F., Lin, D. Y., Lai, T. H., Chiang, H. S. & Lin, Y. H. 2015. SEPT12/SPAG4/LAMINB1 complexes are required for maintaining the integrity of the nuclear envelope in postmeiotic male germ cells. PLos One, 10, e0120722.

Yeh, C. H., Wang, Y. Y., Wee, S. K., Chen, M. F., Chiang, H. S., Kuo, P. L. & Lin, Y. H. 2019. Testis-Specific SEPT12 Expression Affects Sun Protein Localization and is Involved in Mammalian Spermiogenesis. Int J Mol Sci, 20.

Zante, J., Schumann, J., Göhde, W. & Hacker, U. 1977. Dna-fluorometry of mammalian sperm. Histochemistry, 54, 1–7.

