## Supplemental material for "Protamine 2 Deficiency Results In Septin 12 Abnormalities"

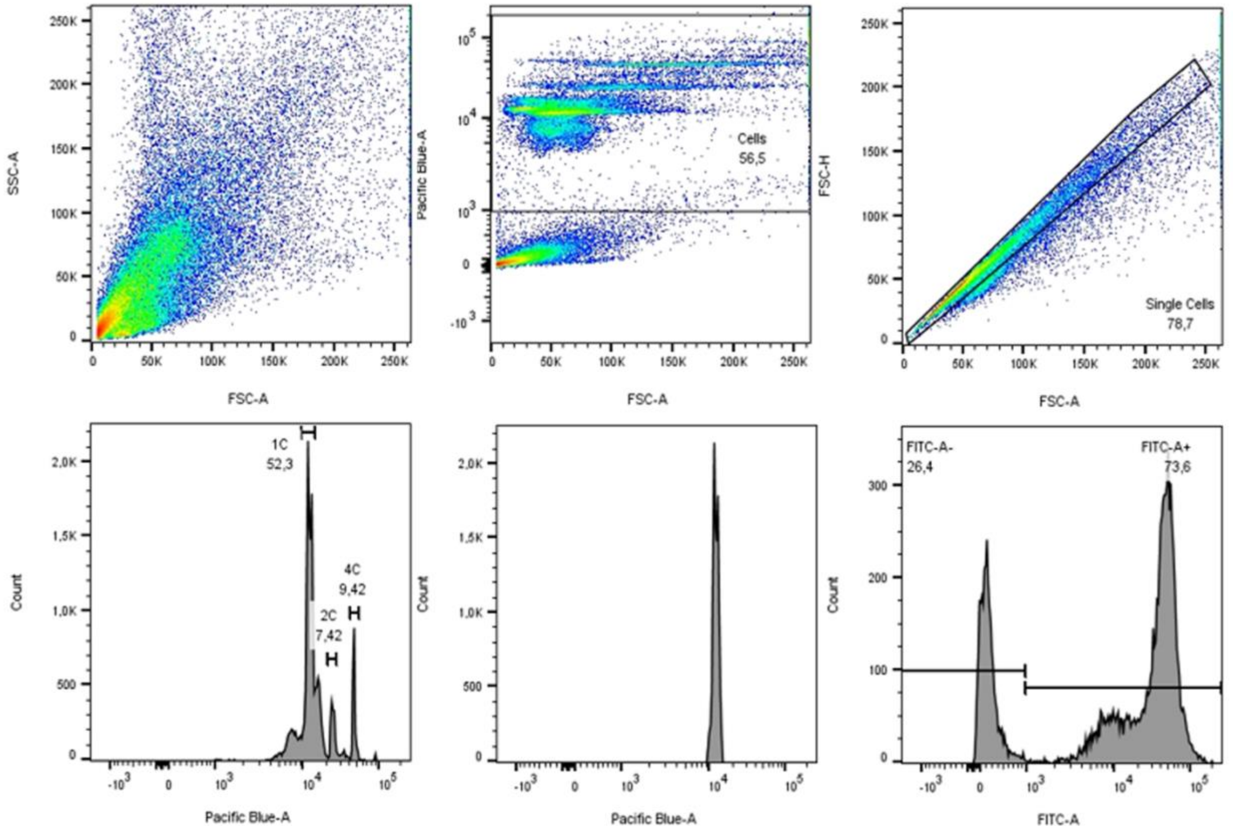


***Supplementary Figure S1: Gating strategy of FCM.*** *Analysis of testicular cell suspension stained with DAPI and antibodies against histone modifications.*

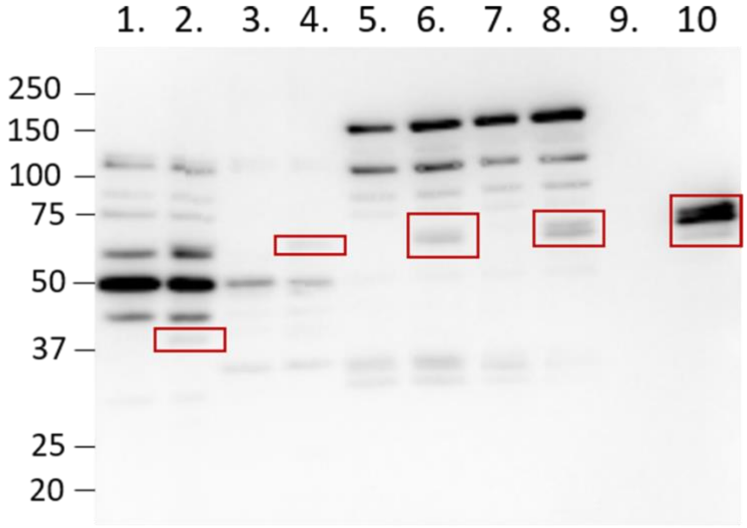


***Supplementary Figure S2: Western blot validation of anti-Septin12 antibody****. Subcellular fractions of non-transfected and transfected HEK293T/17 cells. HEK93T/17 cells were transfected with mouse Septin12-GFP plasmid, red boxes mark positive signals. 1 Non-transfected cells cytoplasmic proteins; 2 Septin 12 Transfected cells cytoplasmic proteins; 3 Non-transfected cells membrane proteins; 4 Septin12 transfected cells membrane proteins; 5 Non-transfected cells nuclear protein; 6 Septin12 transfected cells nuclear protein, 7 Non-transfected cells chromatin-bound nuclear proteins; 8 Septin12 transfected cells chromatin-bound nuclear proteins; 9 Non-transfected cells Tubulin/mitochondria protein fraction; 10 Septin12 transfected cells Tubulin/mitochondria protein fraction.*

| | WT (n=51) | Prm2$^{-/-}$ (n=76) |
|---|---|---|
| Mean signal intensity | 15.92 ± 6.46 | 11.00 ± 3.13 |
| Maximum signal intensity | 51.36 ± 21.07 | 33.62 ± 10.22 |
| Average length | 0.47 ± 0.09 µm | 0.48 ± 0.10 µm |
| Average width | 0.74 ± 0.20 µm | 0.81 ± 0.15 µm |

***Supplementary Figure S3: CEP135 parameters**. Mean and maximum signal intensity of CEP135 staining and average length and width of centrioles in WT and Prm2$^{-/-}$ caudal sperm.*

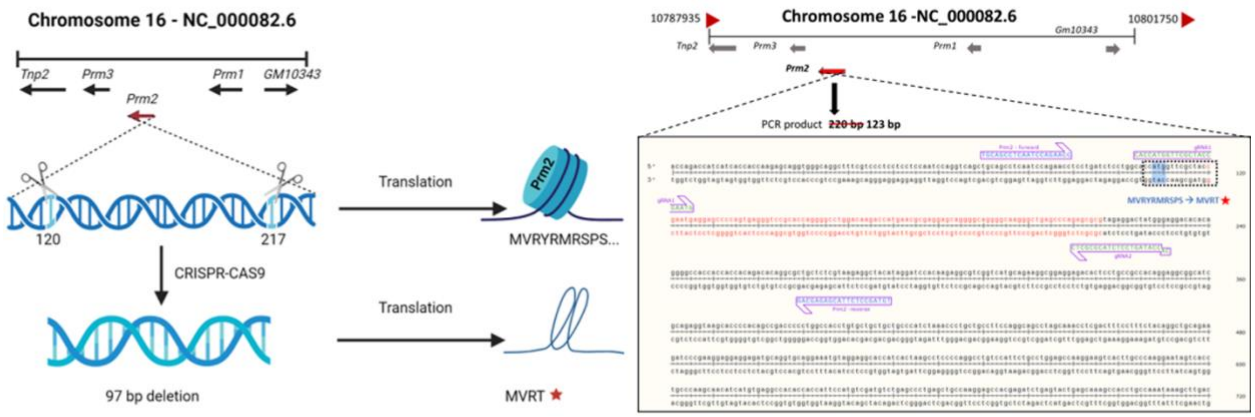


***Supplementary Figure S4: Graphical description of mouse model used in this study**. **(A)** 97 bd deletion was introduced in exon 1 of Prm2 gene, resulting in premature stop codon occurrence. **(B)** Annotation of the knockout mouse line based on the sequencing. The deleted sequence is in red. gRNA sequences used in CRISPR-CAS9 are shown in green, primer sequences used for genotypization are shown in blue. Asterisk mark stop codon.*
